# Dissecting gating mechanisms of Orai calcium channel paralogs using constitutively active Orai mutants that mimic STIM1-gated state

**DOI:** 10.1101/2021.10.26.465861

**Authors:** Bartłomiej Augustynek, Gergely Gyimesi, Jan Dernič, Matthias Sallinger, Giuseppe Albano, Gabriel J. Klesse, Palanivel Kandasamy, Herwig Grabmayr, Irene Frischauf, Daniel G. Fuster, Christine Peinelt, Matthias A. Hediger, Rajesh Bhardwaj

## Abstract

In humans, there are three paralogs of the Orai Ca^2+^ channel, which lie at the heart of the store-operated calcium entry (SOCE) machinery. While the STIM-mediated gating mechanism of Orai channels is still being actively investigated, several artificial and natural variants are known to cause constitutive activity of the human Orai1 channel. Surprisingly, little is known about the conservation of the gating mechanism among the different human Orai paralogs and orthologs in other species. In our work, we show that the mutation corresponding to the activating mutation H134A in transmembrane helix 2 (TM2) of human Orai1 also activates Orai2 and Orai3, likely via a similar mechanism. However, this cross-paralog conservation does not apply to the “ANSGA” nexus mutations in TM4 of human Orai1 which mimic the STIM1-activated state of the channel. Investigating the mechanistic background of these differences, we identified two positions, H171 and F246 in human Orai1, which directly control the channel activation triggered by the “ANSGA” mutations in Orai1. Our results shed new light on these important gating checkpoints and show that the gating mechanism of the Orai channels is affected by multiple factors that are not necessarily evolutionarily conserved, such as the TM4-TM3 coupling.

## Introduction

Store-operated calcium entry (SOCE) is a ubiquitous mechanism by which non-excitable cells regulate basal cytosolic calcium levels and the replenishment of intracellular Ca^2+^ stores. This mechanism is critically important as calcium participates in many different signaling pathways that play central roles in a wide range of cellular processes such as cell division, growth, differentiation, metabolism, gene expression, immune function and others. The SOCE machinery is a multi-component system in which the key components known as Orai proteins are present in the plasma membrane, where they form the pores of the Ca^2+^ release-activated Ca^2+^ (CRAC) channels (Feske et al., 2006; Prakriya et al., 2006; Vig et al., 2006), and the Ca^2+^-sensing STIM proteins are anchored in the ER membrane, from where they regulate gating and activity of the Orai channels (Liou et al., 2005; Roos et al., 2005). In humans, there are three known Orai paralogs (Orai1-3; referred to as hO1, hO2 and hO3, respectively) and two STIM paralogs (STIM1 and STIM2). Furthermore, due to the tissue-specific mRNA splicing, several splice variants of these proteins have been identified (Berna-Erro, Jardin, Salido, & Rosado, 2017; Darbellay, Arnaudeau, Bader, Konig, & Bernheim, 2011; Fukushima, Tomita, Janoshazi, & Putney, 2012; Knapp et al., 2020; Miederer et al., 2015; Niemeyer, 2016; Ramesh et al., 2021; Rana et al., 2015).

Several structural studies have shown that the CRAC channel pore comprises a hexameric arrangement of Orai proteins (Hou, Burstein, & Long, 2018; Hou, Outhwaite, Pedi, & Long, 2020; Hou, Pedi, Diver, & Long, 2012; Liu et al., 2019). The Orai protein itself has four transmembrane (TM) helices, which are arranged radially around the central pore that is formed out of six TM1 helices, one from each subunit (Figure 1). Additional twelve TM helices (6x TM2 and 6x TM3) are arranged in a second, interwoven ring wrapping around the pore-forming TM1 helices. Finally, six TM4 helices form a third ring at the distal region (Hou et al., 2012). Interfaces between the four helices TM1-4 show tight packing of amino acid sidechains. The N-terminus of TM1 that extends into the cytoplasm with residues of unresolved structure encompasses a region (residues 39-59) dispensable for SOCE but critical for ensuring coupling of hO1-mediated local Ca^2+^ entry to the activation of NFAT1 (Nuclear Factor of Activated T-cells) transcription factor (Kar et al., 2021). Notably, the TM4 features a kink at a highly conserved transmembrane proline residue (Pro245 in hO1), followed by another hinge region with the conserved cytosolic sequence of LVSHK (L261-K265 in hO1) residues, also called the “nexus” region (Y. Zhou et al., 2016), and finally a C-terminal, cytosolic extension helix (TM4ext). The first structural studies based on *Drosophila melanogaster* Orai (dOrai) described TM4ext as being locked in a so-called “latched” state, nearly parallel to the membrane bilayer by coiled coil interactions of pairs of antiparallel TM4ext helices between neighboring Orai subunits (Hou et al., 2012). Whether the latched state is a true representation of the native quiescent state of the Orai1 channel is contested and is discussed later including insights from new structural findings.

**Figure 1.**
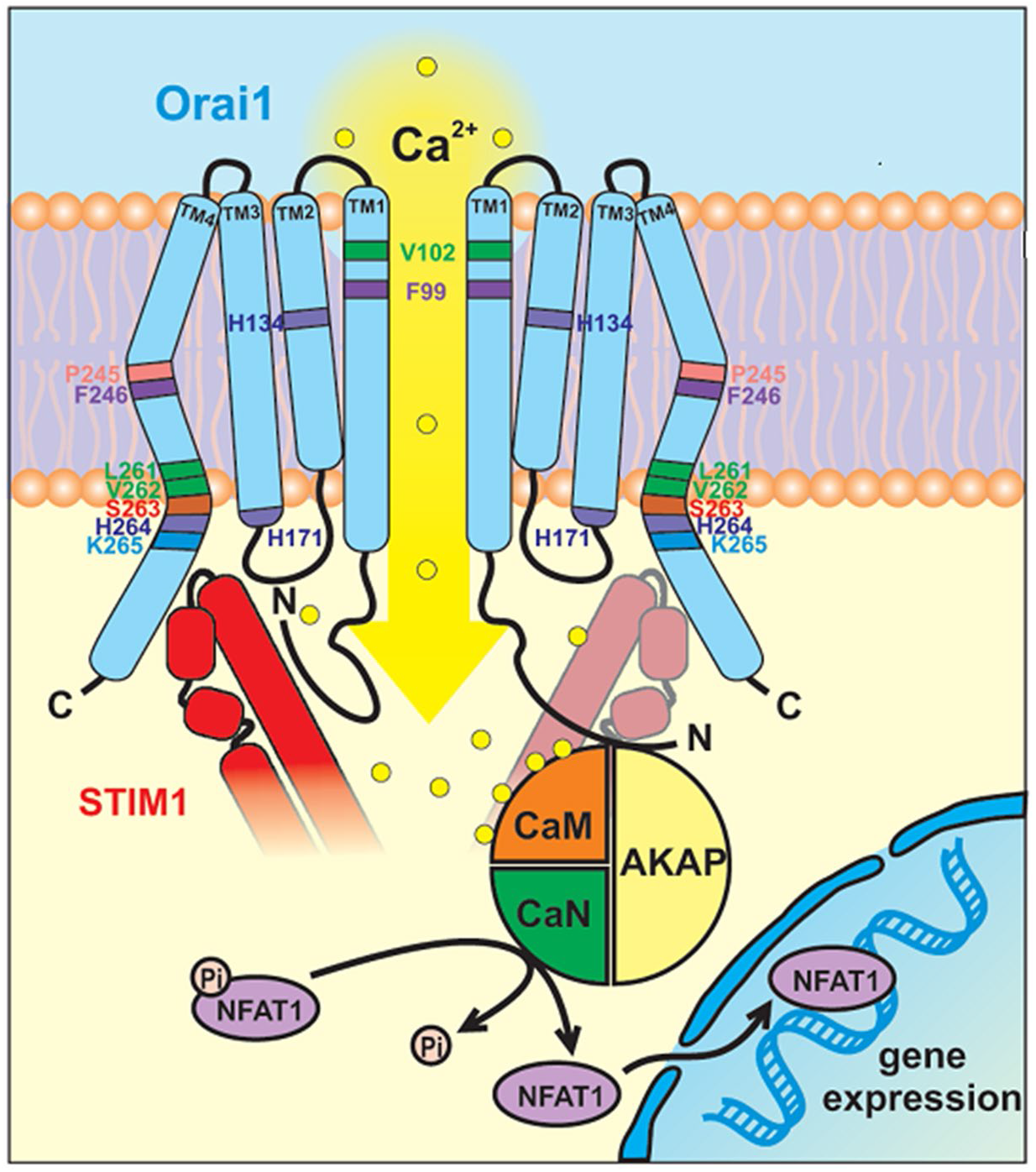
Cartoon representation of human Orai1 showing its four transmembrane (TM) helices TM1-4. Only two out of six subunits are shown for simplicity, highlighting the hydrophobic gate residues F99 and V102 in the TM1 pore helix, H134 in the TM2, H171 in TM3, P245 and F246 in TM4 and LVHSK residues in the TM4 extension (TM4ext). Binding of STIM1/SOAR (shown in red) to the C-terminus of Orai1 leads to conformational changes within Orai1 propagating from TM4 to TM1 leading to opening of the channel pore and influx of Ca2+. The association of A-kinase anchoring protein (AKAP79) accommodating calmodulin (CaM) and calcineurin (CaN) with the N-terminus of Orai1 delimits the dephosphorylation of NFAT1 at the mouth of the channel, and thus facilitating its rapid translocation to the nucleus, wherein it modifies expression of target genes.

Mutations in several key regions of the Orai proteins have shown constitutive activity independent of STIM1 gating. These variants have been instrumental to understand the gating mechanism of Orai channels (Krizova, Maltan, & Derler, 2019; Yeung, Yamashita, & Prakriya, 2020). The hO1 pore-lining residues F99 and V102 constitute the hydrophobic gate of the CRAC channel (McNally, Somasundaram, Yamashita, & Prakriya, 2012; Yamashita et al., 2017) (Figure 1). Pore mutations in TM1, such as V102C, cause constitutive channel opening by physically removing gating barriers within the pore (McNally et al., 2012; Yamashita et al., 2017). However, several gain-of-function mutations are known that are not located in the pore, but in the second (TM2-TM3) or third ring (TM4) of interacting transmembrane helices. These mutant variants are interesting because they can potentially trigger gating-related conformational changes that are downstream to STIM1-based activation (Krizova et al., 2019; Yeung, Yamashita, et al., 2020). Unexpectedly, mutation of residue H134 in hO1, which is located in the TM2 region (second ring of TM helices), constitutively activates hO1 as well as *Drosophila melanogaster* Orai (dOrai) and retains all hallmark properties of CRAC channel activity, such as Ca^2+^ selectivity (Frischauf et al., 2017; Hou et al., 2018; Yeung et al., 2018). The H134 position in hO1 (H206 in dOrai) seems to constitute a “steric brake”, which, when replaced by a residue with a smaller sidechain, causes a significant dilation of the pore (Frischauf et al., 2017; Hou et al., 2018; Hou et al., 2020; Yeung et al., 2018) and “rotation” of the TM1 helix (Bulla et al., 2019; Frischauf et al., 2017; Hou et al., 2018; Yeung et al., 2018). The latter in turn, causes a rearrangement of the pore-lining hydrophobic gate, formed by residues F99 and V102, which is measured as the rotation of the F99 phenyl ring relative to the axis of the pore (Yeung et al., 2018). Opening of the pore via the rearrangement of F99 seems to be mediated by a so-called sulfur-aromatic latch, where M101 of a given hO1 subunit contacts F99 of the neighboring hO1 subunit, stabilizing the open channel conformation. Conversely, in the closed state, M101 forms contacts with F187 (on TM3), highlighting the role of TM1-TM3 crosstalk in channel gating (Bonhenry, Schober, & Schindl, 2021; Yeung, Ing, Yamashita, Pomes, & Prakriya, 2020). Other TM1-TM3 contacts towards the cytoplasmic side have also been shown to be essential for proper channel gating (Dong et al., 2019; Liu et al., 2019; Y. Zhou et al., 2016). On the other hand, the rotation of the TM1 helix was not observed in the molecular dynamics (MD) simulations of the hO1-H134A homology model (Frischauf et al., 2017). Further, the TM1 helix rotation was also not apparent from a recent 3.3 Å resolution cryo-EM structure of dOrai-H206A (corresponding to H134 in hOrai1). However, the displacement of F171 (corresponding to hO1 F99) away from the pore was observed, which likely resulted from the rigid-body outward movement of each dOrai-H206A subunit (Hou et al., 2020). Overall, a common consensus among these findings suggests that sustained Orai1 channel opening requires the displacement of F99 away from the pore.

More enigmatic is the mechanism of how mutations in the third ring (TM4) lead to constitutive channel opening. The notorious P245L mutation of hO1 was found as a gain-of-function mutation in patients with tubular aggregate myopathy (TAM) (Liu et al., 2019; Nesin et al., 2014), rendering the hO1 channel constitutively active, albeit with a significant loss of selectivity towards Ca^2+^ (Liu et al., 2019; Palty, Stanley, & Isacoff, 2015). Structure of P288L dOrai (corresponding to P245L hO1) shows a clear straightening of the kink in TM4 at P288 (Liu et al., 2019). Nevertheless, the relevance of this conformational change has been contested due to establishment of extensive artificial contacts between Orai hexamers, present only in protein crystals but not in physiological conditions, and to the fact that it shows an apparently closed pore geometry (Hou et al., 2020). Interestingly, a non-natural mutant variant of TM4 in hO1, in which the “LVSHK” hinge sequence is replaced by “ANSGA”, reproduces all hallmark properties of CRAC current (*I*_CRAC_), including inward rectification and Ca^2+^-selectivity. This contrasts with the P245L variant (Y. Zhou et al., 2016). Thus, the ANSGA substitution, being closest to the STIM1-binding site in TM4ext (Frischauf et al., 2009; Z. Li et al., 2011; Muik et al., 2008; Navarro-Borelly et al., 2008; Palty et al., 2015; Park et al., 2009; Tirado-Lee, Yamashita, & Prakriya, 2015; Y. Zhou et al., 2016), makes this variant a potentially useful tool to study STIM1-like activation of Orai channels. However, currently no structural information is available about the ANSGA variant and its gating mechanism.

One of the interesting features of the X-ray-based structures of dOrai is that the TM4ext helices protrude either in a “latched” (“quiescent”) (Hou et al., 2012) or an “unlatched” state, where the latter state displays the dissociation of the TM4ext bundles, a clear straightening of the kinks in TM4 at both the P288 and the nexus position, and rupture of contacts between TM3 and TM4 (Hou et al., 2018; Liu et al., 2019; Y. Zhou et al., 2019). Importantly, it was concluded that the latched state prevents pore opening, and unlatching is a necessary, but not sufficient condition of channel opening (Hou et al., 2018). However, the physiological relevance of these TM4ext bundles has recently been disputed, as they might be artificially stabilized by crystal contacts, and therefore the role of unlatching in channel activation and pore opening became less clear (Y. Zhou et al., 2019). Further, the interactions between plasma membrane targeted C-terminal TM4ext peptides of hO1 or hO3 could not be observed in a recent FRET-based study (Baraniak et al., 2021) which further questions the formation of antiparallel TM4ext interaction pairs between neighboring Orai subunits as reported in the closed dOrai structure (Hou et al., 2012). Future structural studies addressing the closed state of the Orai channel should settle this debate. In context of the channel activation pathway, in particular, TM3-TM4 contacts were proposed to be integral parts (Liu et al., 2019; Y. Zhou et al., 2016; Y. Zhou et al., 2019), since cross-linking studies of positions L174-L261 showed that TM3 and TM4 can also form contacts in the STIM1-activated state, leading to further enhancement of the CRAC current (Y. Zhou et al., 2016). Consistently, disruption of a cluster of hydrophobic residues between TM3 and TM4 (L174, F178 in TM3 and L261, F257 in TM4) attenuates channel activation (Liu et al., 2019; Y. Zhou et al., 2016). Indeed, cryo-EM studies, which avoid the formation of crystal contacts, have reported an apparently semi-unlatched state of dOrai P288L (Dong et al., 2019; Liu et al., 2019; Y. Zhou et al., 2019), and also an open state of H206A mutant with intact TM3-TM4 contacts (Hou et al., 2020). It is important to mention that while the relevance of TM4ext latching and unlatching is still debated, dissociation of TM4ext bundles is expected since known interaction sites of Orai1 with STIM1 (L273 and L276) lie in the coiled-coil TM4ext region of Orai (Frischauf et al., 2009; Z. Li et al., 2011; Muik et al., 2008; Navarro-Borelly et al., 2008; Palty et al., 2015; Tirado-Lee et al., 2015). Nevertheless, it is intriguing to speculate whether and how the ANSGA substitution may interfere with the TM4ext regions or TM3-TM4 contacts. The contacts between cytoplasmic regions of TM1 and TM3 may also be essential for the constitutive activity of hO1-ANSGA channel (Y. Zhou et al., 2016). In fact, the K85E and L174D hO1 mutations that abrogate *I*_CRAC_ (Lis, Zierler, Peinelt, Fleig, & Penner, 2010; McNally, Somasundaram, Jairaman, Yamashita, & Prakriya, 2013; Y. Zhou et al., 2016) likely by disrupting the TM1-TM3 and TM3-TM4 cytoplasmic contacts, respectively (Dong et al., 2019; Liu et al., 2019; Y. Zhou et al., 2016), not only abolished the constitutive activity of ANSGA but also that of the P245L mutant channel (Derler et al., 2018; Y. Zhou et al., 2016). It remains puzzling why hO1-ANSGA shows CRAC-like activation while P245L does not, despite their proximity and their location on TM4, indicating that factors other than unlatching or TM4 rearrangement are also affecting the selectivity of ion permeation through the Orai hexamer.

Although all three hOrai paralogs undergo STIM1-mediated activation of CRAC currents, hO2 and hO3 generate 2-3 fold lower currents compared to hO1 (Frischauf et al., 2009; Lis et al., 2007; Mercer et al., 2006). Certain regions of the Orai channels are responsible for paralog-specific differences in gating by STIM1 (Fahrner et al., 2018). Deletion of more than 74 residues from the N-terminus of hO1 completely abolishes STIM1-gated CRAC currents and results in significantly attenuated association of hO1 with STIM1 (Derler et al., 2013; Z. Li et al., 2007; Lis et al., 2010; McNally et al., 2013; Muik et al., 2008; Park et al., 2009; Zheng et al., 2013). Interestingly, N-terminal deletion mutants of hO1 can be rescued by replacing loop2 of hO1 (the cytoplasmic region between TM2 and TM3) with that of hO3 (Fahrner et al., 2018). This is surprising given the fact that loop2 is very conserved among Orai paralogs, and the effect of loop2 replacement can be narrowed down to the replacement of 5 amino acids from hO1 with the corresponding ones from hO3 (N147H/K161H/E162Q/E166Q/H171Y), reproducing the rescue effect (Fahrner et al., 2018). These results indicate an extensive crosstalk between TM1 and loop2, which, considering the role of TM1-TM3 contacts in the transduction of gating signals, likely shape paralog-specific differences in gating among human Orai channels. Another recent study elucidates the paralog-specific function of hO1 *vs*. hO3 gating checkpoints in TM3 (Tiffner et al., 2021). Additional paralog-specific differences in gating of human Orais stem from the differential interaction of their C-termini with STIM1 (Alansary, Bogeski, & Niemeyer, 2015; Baraniak et al., 2021; Frischauf et al., 2009; S. Li et al., 2019; Niu et al., 2020). Despite these differences in gating between Orai paralogs, the effects of mutations causing constitutive activity mimicking a STIM1-bound state on various Orai channels have not been studied in detail.

The main objective of the current study was to generate new insights into the gating mechanisms of Orai channels and to investigate if there are paralog-specific differences in conformational rearrangements that are linked to Orai channel gating. To address this, we used the H134A and ANSGA hO1 mutants that are reported to largely mimic the STIM1-gated state of the Orai1 channel (Frischauf et al., 2017; Yeung et al., 2018; Y. Zhou et al., 2016). We constructed their corresponding hO2 and hO3 variants and assessed their constitutive activity by Ca^2+^ imaging on Fluorometric Imaging Plate Reader (FLIPR), electrophysiology as well as NFAT1 nuclear translocation assays. Herein, we show that the conformational coupling between TM2 and TM1 is conserved among all three human Orai paralogs. We investigated whether and how, based on their location, the ANSGA and other residues in Orai paralogs adjacent to the ANSGA sequence might interfere with the TM4ext regions or TM3-TM4 contacts, and thus with the channel activation *per se*. Our data reveal that the TM4-TM3 coupling implicated in the activation of hO1 likely works differently for the paralogs hO2 and hO3. Overall, our data provides novel insights into the gating mechanisms of Orai channels.

## Results

### The human Orai1 TM2 H134A as well as TM3 F187C corresponding mutations in Orai2 and Orai3 renders them constitutively active

The hO1 TM2 residue H134 (**Figure 2A**), which makes the channel constitutively active when mutated to alanine by displacing the TM1-F99 residue away from the pore axis, is highly conserved across multiple species (mouse, *Xenopus* and *Drosophila*) as well as in the hO2 and hO3 paralogs (**Figure 2B**). In order to investigate whether the H134A corresponding mutation in hO2 and hO3 mimics the constitutive activation effect, we generated hO2-H108A and hO3-H109A mutants and expressed them in STIM1/STIM2 double knock-out (S1/S2 DKO) HEK293 cells for electrophysiological analysis. The S1/S2 DKO HEK293 cells were generated in-house using the CRISPR/Cas9 technique specifically to exclude the canonical STIM-dependent activation of Orai channels. The successful ablation of S1 and S2 proteins was confirmed at mRNA and protein levels (**Figure 2—figure supplement 1A,B**). As expected, SOCE was nearly completely absent in the DKO cells (**Figure 2—figure supplement 1C,D**). When expressed in these cells, the hO2-H108A and hO3-H109A mutants showed constitutive currents similar to hO1-H134A, which disappeared upon removal of Ca^2+^ from the bath solution, whereas the wild type (WT) Orai channels did not show any constitutive activity (**Figure 2C**). Consistent with this finding, nearly 90% of HEK293 cells expressing either hO1-H134A, hO2-H108A or hO3-H109A mutant displayed translocation of the CFP labeled NFAT1 transcription factor from cytosol to nuclei, whereas only less than 10% of the cells expressing WT Orai proteins showed nuclear translocation of NFAT1 (**Figure 2D,E**). Overall, these results suggest that the substitution of H108 of hO2 and H109 of hO3 by alanine exerts a similar constitutive activation effect as known for the hO1-H134A mutant.

**Figure 2.**
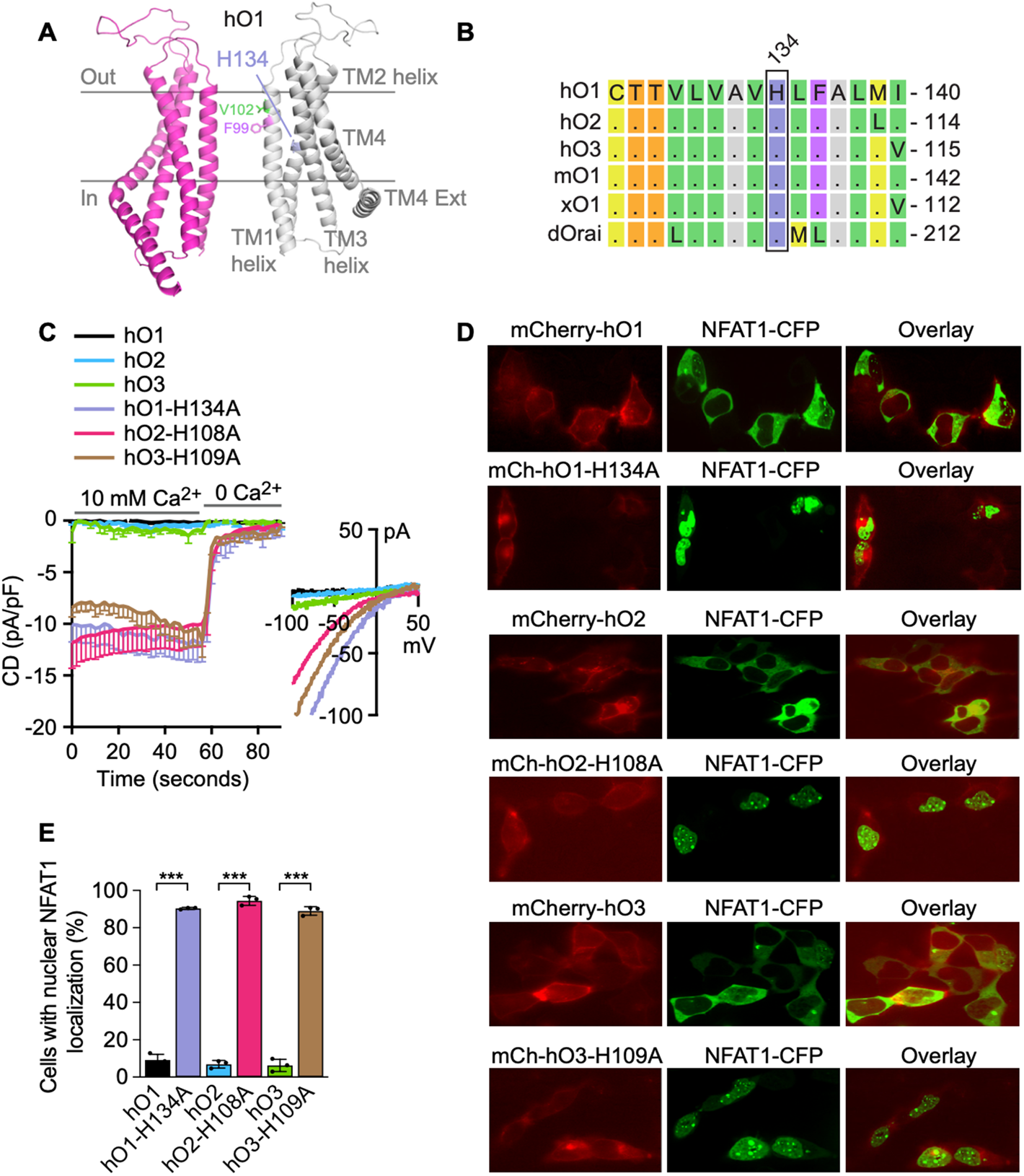
TM2 Histidine mutations lead to constitutive Ca2+ influx through human Orai channels leading to NFAT1 activation. (**A**) Cartoon representation of the homology model of hO1 channel showing only two subunits of the hexameric model based on the *Drosophila* Orai model (PDB ID: 4HKR). TM helices TM1-4 are shown. (**B**) Multiple sequence alignment of TM2 of human Orai1 (hO1), human Orai2 (hO2), human Orai3 (hO3), mouse Orai1 (mO1), *Xenopus laevis* Orai1 (xO1) and *Drosophila melanogaster* Orai (dOrai) is shown highlighting the conserved H134 residue. (**C**) Current densities (CD) of the constitutive Ca2+ currents recorded from HEK293 STIM1/STIM2 (S1/S2) double knockout (DKO) cells transiently overexpressing: WT hO1 (n=6), WT hO2 (n=9), WT hO3 (n=7), hO1-H134A (n=6), hO2-H108A (n=10) and hO3-H109A (n=6), presented as average values, -SEM, with corresponding average current-voltage (I/V) relations extracted at t= 59s. (**D**) Representative fluorescence and confocal microscopy images of HEK293 cells co-expressing NFAT1-CFP with either of the indicated mCherry-Orai constructs along with the CFP/mCherry overlay images. (**E**) HEK293 cells expressing indicated constructs with nuclear NFAT1-CFP localization shown as percentage (mean ±SD; n=3). P value (*p*) ≤ 0.001 is indicated as “***”.

Recently, it has been proposed that intersubunit F99-M101 interactions support the hO1 channel opening, whereas intersubunit M101-F187 (TM1-TM3) interactions facilitate the closing of the channel (Bonhenry et al., 2021; Yeung, Ing, et al., 2020). Our investigations of these interactions using MD simulations revealed that H134S/H134A hO1 open channel mutants have a higher frequency of intersubunit F99-M101 contacts than WT, whereas no difference was observed in M101-F187 interactions (**Figure 2—figure supplement 2C,D,E**). These observations are in line with the recently published findings on dOrai-H206Q/C channels (Yeung, Ing, et al., 2020). In view of the role of M101-F187 interactions in maintaining channel closure (Yeung, Ing, et al., 2020); using the FLIPR-based Ca^2+^ imaging technique, we first validated the earlier published finding that F187C substitution in hO1 results in its constitutive activation (Yeung et al., 2018) (**Figure 2—figure supplement 2F,G**). Next, given the conservation of M101 and F187 residues in hO2 and hO3 (**Figure 2—figure supplement 2A,B**), we investigated if the mutations corresponding to hO1 F187C, F161C and F162C in hO2 and hO3, respectively, also lead to their constitutive activation. Our results showing the constitutive activation of hO2-F161C and hO3-F162C (**Figure 2—figure supplement 2H,I**) suggest that intersubunit TM1-TM3 (M101-F187) interactions facilitating the closed state of the hO1 channel (Bonhenry et al., 2021; Yeung, Ing, et al., 2020) may also be conserved in hO2 and hO3.

### The substitution of Orai1 nexus LVSHK residues with ANSGA results in constitutive activation of human and *Xenopus* Orai1 but not human Orai2, Orai3 and *Drosophila* Orai

Next, we investigated the only other known Ca^2+^-selective, constitutively active nexus mutant of hO1 where the LVSHK residues (positions 261-265 in hO1), forming a nexus connecting TM4 with its cytoplasmic C-terminal extension, are mutated to ANSGA residues (**Figure 3A**). The LVSHK nexus motif is completely conserved in hO1, mouse Orai1 (mO1), *Xenopus laevis* Orai1 (xO1) and dOrai, whereas the serine at 263 residue position in corresponding hO2 and hO3 sequences is substituted with arginine and alanine, respectively (**Figure 3B**). The hO1-ANSGA nexus mutant expressed in S1/S2 DKO cells, as expected, exhibited constitutive influx of Ca^2+^ in both Ca^2+^ imaging (**Figure 3C,D**) and patch-clamp measurements (**Figure 3G**). Strikingly, analogous experiments with the corresponding hO2-ANSGA or hO3-ANSGA variants revealed no constitutive entry of Ca^2+^ through these mutants (**Figure 3E,F,G**). Consistently, the hO2-ANGSA or hO3-ANSGA expression did not result in any significant increase in the percentage of HEK293 cells with nuclear translocation of NFAT1 compared to the expression of their WT counterparts (**Figure 3J,K)**. However, nearly 90% of cells expressing hO1-ANSGA showed nuclear translocation of NFAT1 compared to less than 10% for WT hO1 (**Figure 3J,K).** Next, we investigated whether introduction of the ANSGA mutation in the nexus region of hO1 orthologues found in lower organisms can render them constitutively active. Despite the conserved LVSHK motif in both xO1 and dOrai, only the xO1-ANSGA mutant exhibited constitutive Ca^2+^ entry (**Figure 3H**) while dOrai remained inactive similarly to its WT counterpart (**Figure 3I**). In order to rule out the possibility that introduction of the ANSGA mutation compromised surface expression of hO2-ANSGA, hO3-ANSGA and dOrai-ANSGA, we introduced an additional, well-established mutation (corresponding to V102A pore-opening mutation of hO1) directly into the TM1 of these mutants. Indeed, the introduction of V76A, V77A and V174A in hO2-ANSGA, hO3-ANSGA and dOrai-ANSGA, respectively, resulted in constitutive Ca^2+^ influx through these mutants, suggesting that ANSGA mutation does not affect their cell surface localization or proper assembly (**Figure 3—figure supplement 1A,B,C**). Altogether, these data establish that mutation of nexus residues to ANSGA does not lead to constitutive activation of hO2, hO3 and dOrai, but constitutively activates hO1 and xO1.

**Figure 3.**
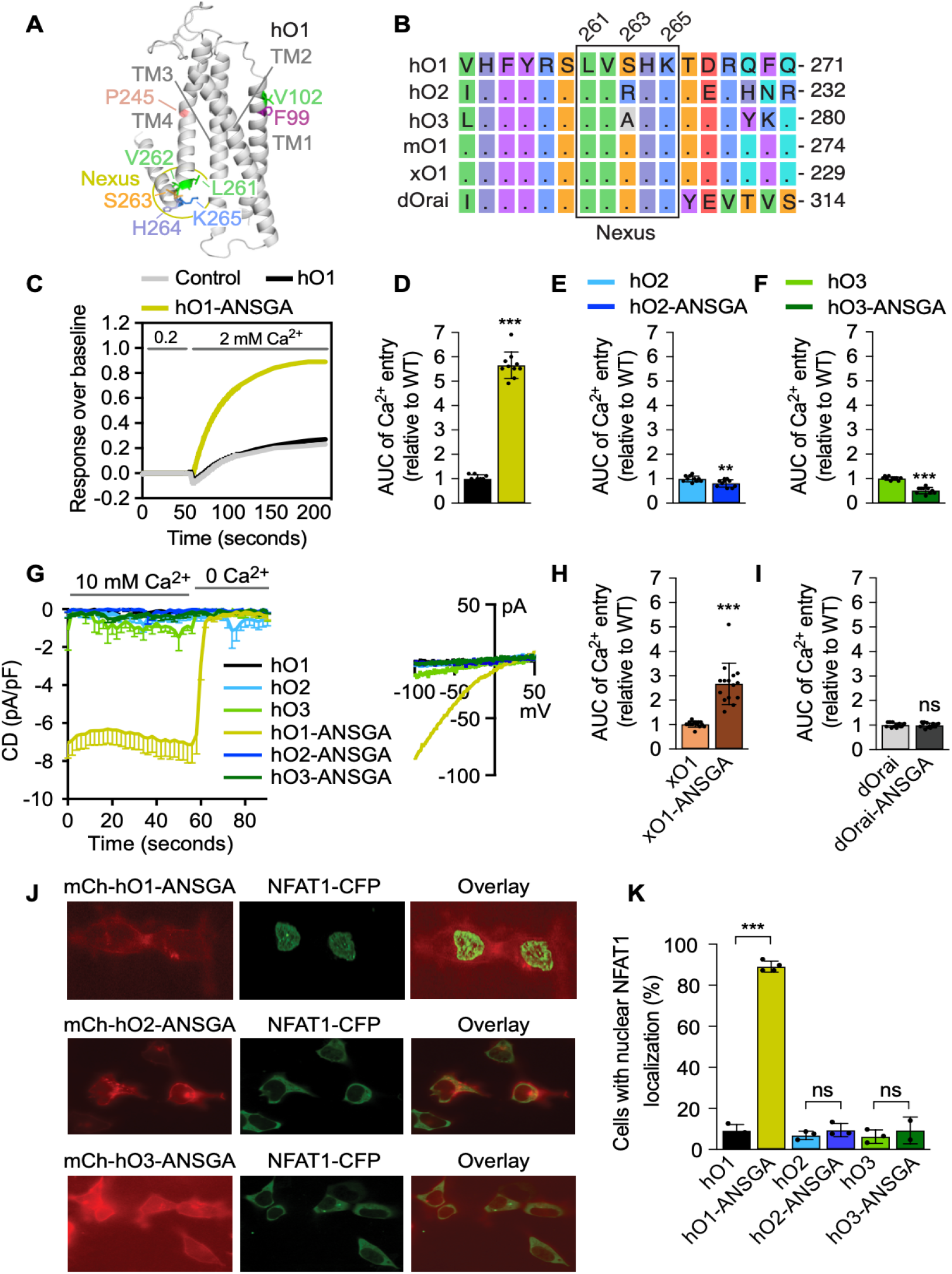
ANSGA substitution of the Orai nexus residues does not exert a constitutive activation effect among Orai homologs. (**A**) Cartoon representation of a single subunit of hO1 homology model depicting indicated residues along with the LVSHK (261-265) nexus residues. (**B**) Multiple sequence alignment of TM3 of hO1, hO2, hO3, mO1, xO1 and dOrai is shown, highlighting the conserved LVXHK residues. (**C**) Representative constitutive Ca^2+^ entry traces recorded from HEK293 S1/S2 DKO cells transfected with mCherry control vector, WT hO1 and hO1-ANSGA constructs with initial baseline recording in 0.2 mM CaCl2, followed by addition of 2 mM CaCl2. The quantified AUC of Ca^2+^ entry peak from HEK293 S1/S2 DKO cells expressing (**D**) WT hO1 and hO1-ANSGA, (**E**) WT hO2 and hO2-ANSGA and (**F**) WT hO3 and hO3-ANSGA (mean ± SD; n=10). *p* ≤ 0.001 is indicated as “***” and 0.001 < *p* ≤ 0.01 is indicated as “**”. (**G**) Current densities (CD) of the constitutive Ca^2+^ currents recorded from HEK293 S1/S2 DKO cells transiently overexpressing: WT hO1 (n=6), WT hO2 (n=9), WT hO3 (n=7), hO1-ANSGA (n=23), hO2-ANSGA (n=6) and hO3-ANSGA (n=7), presented as average values, -SEM, with corresponding average current-voltage (I/V) relations extracted at t= 59s. (**H**) AUC of the constitutive Ca^2+^ entry traces from WT xO1 and xO1-ANSGA expressing HEK293 cells (mean ± SD; n=15). *p* ≤ 0.001 is indicated as “***”. (**I**) AUC of the constitutive Ca^2+^ entry traces from S1/S2 DKO HEK293 cells expressing WT dOrai and dOrai-ANSGA constructs (mean ± SD; n=15). *P* ≥ 0.05 is indicated as “ns”. (**J**) Representative confocal microscopy images of HEK293 cells co-expressing NFAT1-CFP with either of the indicated mCherry-Orai-ANSGA constructs along with the CFP/mCherry overlay images. (**K**) HEK293 cells expressing indicated constructs with nuclear NFAT1-CFP localization shown as percentage (mean ± SD; n ≥ 3). *p* ≤ 0.001 is indicated as “***” and *p* ≥ 0.05 is indicated as “ns”.

### The H171Y/F, but not H171A substitution in the TM3 helix of human Orai1-ANSGA abolishes its constitutive activity

We next targeted the molecular determinants that lead to these paralog-specific differences in ANSGA substitution-mediated constitutive activity of Orai channels. For this purpose, we looked in our homology-based model of hO1 for the conservation of residues that are in close proximity to the nexus. We identified a histidine residue at position 171 of hO1 that directly faces the nexus region and is conserved in mO1 and xO1, but is substituted by a tyrosine residue in hO2, hO3 and dOrai (**Figure 4A,B**). Based on this and in light of the previous finding that ANSGA substitution of nexus constitutively activates hO1 and xO1 (**Figure 3D,H**), but not hO2, hO3 and dOrai (**Figure 3E,F,I**), we have hypothesized that the H171Y mutation could interfere with the constitutive activity of the hO1-ANSGA mutant. Indeed, H171Y substitution resulted in complete attenuation of the constitutive activity of hO1-ANSGA channel (**Figure 4C,D**). To investigate whether mislocalization or altered assembly were the reasons for abolished constitutive activity of the hO1-H171Y-ANSGA channel, we used surface biotinylation experiments and confirmed the expression of hO1-H171Y-ANSGA in the plasma membrane (**Figure 4—figure supplement 1A**). In addition, the introduction of the V102C pore mutation rescued the constitutive activity of the hO1-H171Y-ANSGA channel and ensured the correct assembly of the channel (**Figure 4—figure supplement 1B**). We next tested further substitutions of H171, of which the H171F, but not H171A abolished the constitutive activity of the hO1-ANSGA mutant (**Figure 4E,F**). Consistently, both the H171Y and the H171F but not the H171A substitutions prevented the nuclear translocation of NFAT1 (**Figure 4G, Figure 4—figure supplement 1C**).

**Figure 4.**
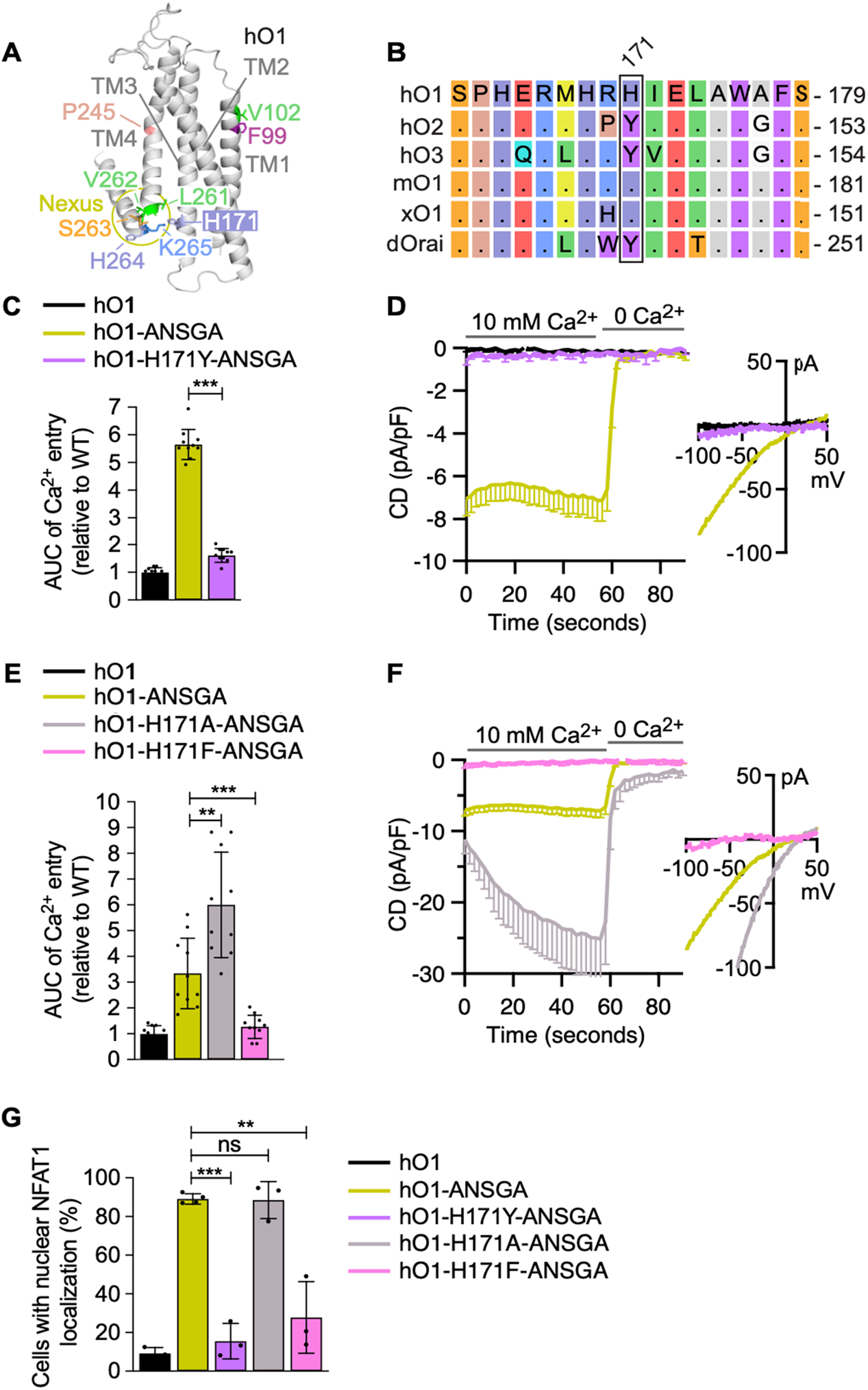
Impact of H171 residue mutations on the constitutive activity of the hO1-ANSGA channel. **(A)** Cartoon representation of a single subunit of hO1 homology model depicting indicated residues along with the LVSHK (261-265) nexus and H171 residues. (**B**) Multiple sequence alignment of a segment of loop2 and TM3 of hO1, hO2, hO3, mO1, xO1 and dOrai is shown, highlighting the residues corresponding to hO1 H171. (**C**) The quantified AUC of constitutive Ca^2+^ entry recorded in HEK293 S1/S2 DKO cells expressing WT hO1, hO1-ANSGA and hO1-H171Y-ANSGA (mean ± SD; n=10). *p* ≤ 0.001 is indicated as “***”. (**D**) Current densities (CD) of the constitutive Ca^2+^ currents recorded from HEK293 S1/S2 DKO cells transiently overexpressing: WT hO1 (n=6), hO1-ANSGA (n=23) and hO1-H171Y-ANSGA (n=8), presented as average values, - SEM, with corresponding average current-voltage (I/V) relations extracted at t= 59s. (**E**) The AUC quantified from constitutive Ca^2+^ entry recordings in HEK293 S1/S2 DKO cells expressing WT hO1, hO1-ANSGA, hO1-H171A-ANSGA and hO1-H171F-ANSGA (mean ± SD; n=10). *p* ≤ 0.001 is indicated as “***” and 0.001 < *p* ≤ 0.01 as “**”. (**F**) Current densities (CD) of the constitutive Ca^2+^ currents recorded from HEK293 S1/S2 DKO cells transiently overexpressing: WT hO1 (n=6), hO1-ANSGA (n=23), hO1-H171A-ANSGA (n=7) and hO1-H171F-ANSGA (n=6), presented as average values, -SEM, with corresponding average current-voltage (I/V) relations extracted at t= 59s. (**G**) HEK293 cells expressing WT hO1, hO1-ANSGA, hO1-H171Y-ANSGA, hO1-H171A-ANSGA and hO1-H171F-ANSGA constructs having NFAT1-CFP localized to the nucleus shown as percentage (mean ±SD; n ≥ 3). P value (*p*) ≤ 0.001 is indicated as “***”, 0.001 < *p* ≤ 0.01 as “**” and *p* ≥ 0.05 indicated as “ns”.

### The H171Y substitution in human Orai1-ANSGA affects the α-helical propensity of the TM4 helix extension as well as the local coupling between TM3 and TM4

We sought to understand the mechanism behind H171Y-mediated inhibition of the constitutive activity of the hO1-ANSGA channel. We hypothesized that the H171Y substitution may disrupt the ANSGA-mediated proposed flexion of the nexus hinge (Y. Zhou et al., 2016) and/or TM4-TM3 coupling (Liu et al., 2019; Y. Zhou et al., 2016; Y. Zhou et al., 2019) within hO1-ANSGA, both of which are considered critical for the constitutive activity of hO1-ANSGA. No difference in the helicity of the TM4 helix region (residues 240-250), encompassing the proline bend caused by P245 was observed in the molecular dynamics (MD) simulations of hO1-ANSGA and hO1 WT (**Figure 5A**, **Figure 5—figure supplement 1A**). Contrariwise, the tubular aggregate myopathy (TAM)-associated P245L mutant showed a significantly higher α-helical propensity of the TM4 fragment upstream of L245 (residues 241-244) compared to WT (**Figure 5—figure supplement 1A**). On the other hand, we observed a modest increase in the α-helical propensity of the nexus hinge upon the substitution of the nexus residues to ANSGA (**Figure 5B**). Further, our MD simulations reveal that H171Y but not H171A substitution led to a decrease in α-helical propensity of residues 265 to 267 of hO1-ANSGA channel (**Figure 5B**), which may contribute to the loss of constitutive activity of the hO1-H171Y mutant ANSGA channel.

**Figure 5.**
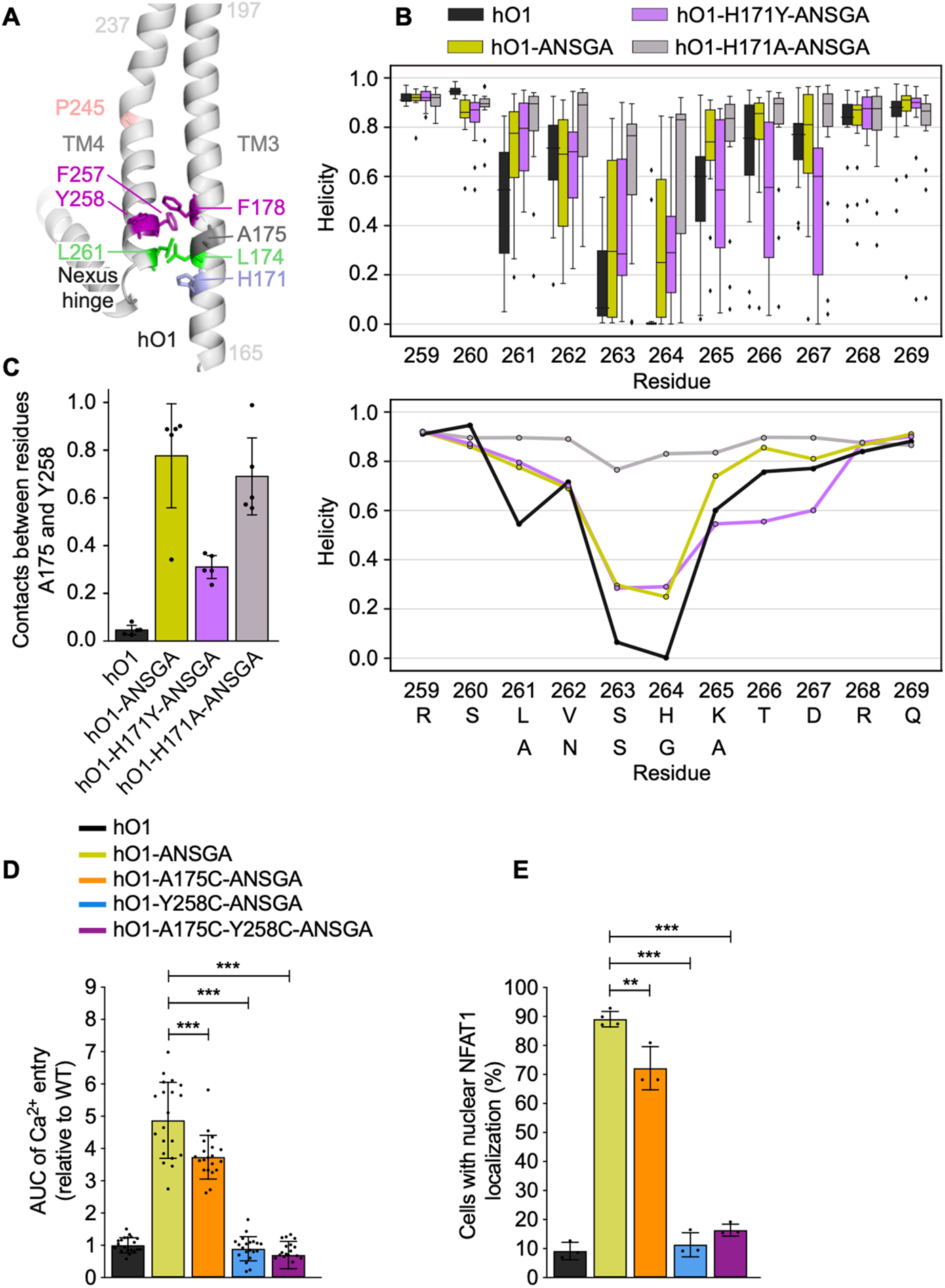
Role of TM3-TM4 contacts and helicity of the nexus segment in the gating of the hO1 ANSGA channel. (**A**) Structural model of hO1 showing the relevant residues in the TM3-TM4 region. (**B**) Helicity of the nexus region averaged over 5 MD trajectories for chains A, C and E, giving 15 points per residue and system (see Methods). In the upper panel, box-and-whiskers plot showing quartiles is used, outliers are plotted individually. In the lower panel, the same data is plotted for clarity using mean values. (**C**) Intra-subunit contact frequencies between residues A175 and Y258 averaged over each of the 5 MD trajectories of WT hO1 and hO1-ANSGA mutant variants. (**D**) The quantified AUC of constitutive Ca2+ entry recordings from HEK293 S1/S2 DKO cells expressing either WT hO1, hO1-ANSGA or indicated mutants (mean ± SD; n=20). *p* ≤ 0.001 is indicated as “***”. (**E**) HEK293 cells expressing indicated mCherry-hO1 or hO1-ANSGA variants with nuclear NFAT1-CFP localization shown as percentage (mean ± SD; n ≥ 3). *p* ≤ 0.001 is indicated as “***” and 0.001 < *p* ≤ 0.01 is indicated as “**”.

Next, we addressed whether ANSGA and further H171Y/A substitutions within hO1-ANSGA affect the frequency of contacts between known, functionally important, interacting residues at the interface of TM3 and TM4, namely, L174(TM3)-L/A261(TM4) and F178(TM3)-F257(TM4) (**Figure 5A**). No differences could be observed in the contact frequencies between these residues in the MD simulations of hO1 WT, hO1-ANSGA, hO1-H171Y-ANSGA and hO1-H171A-ANSGA (**Figure 5—figure supplement 1B**). Interestingly, further screening and contact analysis led to the identification of different contact frequencies between A175 (TM3) and Y258 (TM4). The MD simulations of the hO1-ANSGA channel showed higher A175-Y258 contact frequency compared to the WT channel. Substitution of H171 with tyrosine but not alanine residue resulted in a reduction of A175-Y258 contacts (**Figure 5C**) indicating that the H171Y substitution attenuates the local coupling between TM3 and TM4 in the hO1-ANSGA channel. In line with the previous findings of others, the L174D mutation in the hO1-ANSGA channel abolished its constitutive activity (Y. Zhou et al., 2016). Moreover, both F178A and F257A substitutions, reported to affect the STIM1-mediated gating of the WT Orai1 channel (Liu et al., 2019), diminished the constitutive activity of the hO1-ANSGA channel, further highlighting the importance of these residues in the channel activation (**Figure 5—figure supplement 1C**). Finally, we addressed whether A175 and Y258 are indispensable for the hO1-ANSGA channel activation. While the A175C substitution led to only a small reduction, Y258C resulted in complete loss of constitutive activity of the hO1-ANSGA channel without affecting the cell surface localization (**Figure 5D, Figure 5—figure supplement 1D**). As expected, the A175C-Y258C mutant of the hO1-ANSGA channel did not exhibit constitutive activation. The corresponding effect was observed in NFAT1 activation experiments, wherein expression of hO1-A175C-ANSGA led to only small decrease in percentage of cells showing nuclear translocation of NFAT1 compared to hO1-ANSGA. Conversely, the expression of hO1-Y258C-ANSGA and hO1-A175C-Y258C-ANSGA mutants reduced the number of cells with nuclear NFAT1 to background level, *i.e.*, the cells expressing hO1 WT (**Figure 5E, Figure 5—figure supplement 1E**).

We furthermore investigated whether the pore helix rotation reported for the dOrai mutant corresponding to hO1 H134S (Yeung et al., 2018) or pore dilation reported for human H134A Orai1 mutant (Frischauf et al., 2017), could also be observed in the hO1-ANSGA mutant, and whether the effect of H171Y substitution in hO1-ANSGA channel alters these parameters. As reported earlier, we could observe rotation of the F99 residue indicative of TM1 helix rotation in hO1-H134S MD simulations (Bulla et al., 2019), which was even more evident in hO1-H134A mutant channel (**Figure 5—figure supplement 2A**). However, this effect could neither be observed in constitutively open hO1-P245L TM4 mutant nor in hO1-ANSGA mutant channel in course of 200 ns long simulations (**Figure 5—figure supplement 2A**). On the other hand, in comparison to hO1 WT, no considerable pore dilation was clearly evident in the MD simulations of any of the tested hO1 mutants including hO1-ANSGA (**Figure 5—figure supplement 2B,C**). Since neither TM1 helix rotation nor pore dilation was evident in the MD simulations of hO1-ANSGA, we did not analyze these parameters for the H171 mutants of the hO1-ANSGA channel. Overall, these results indicate that the substitution of H171 by tyrosine in human Orai1-ANSGA decreases the α-helical propensity of the TM4 helix extension (residues 265-267) as well as attenuates the local coupling between TM3 and TM4 (A175-Y258). Additionally, we identified Y258 as a critical residue for the constitutive activity of the hO1-ANSGA mutant channel.

### The H171Y substitution does not abolish the constitutive activity of human Orai1 TM1 V102C, TM2 H134A and TM4 P245L and F250C mutants

Next, we aimed to determine if H171Y mutation also abolishes the activity of other known constitutively open Orai1 mutants, or if its effect is specific to the ANSGA-dependent constitutive activity of hO1. The hO1 TM4 mutants, P245L or F250C as well as TM2 mutant H134A, did not show any deficit in constitutive Ca^2+^ influx when additional H171Y mutation was introduced in these channels. Also, the TM1 pore mutant hO1-V102C upon introduction of H171Y substitution resulted in only a minor drop of constitutive activity (**Figure 6A,B**) suggesting the inhibitory effect of H171Y is specific to hO1-ANSGA. Further, the effect of H171Y on the constitutive activity of the hO1-ANSGA mutant channel was independent of the TM4 cytosolic extension beyond the ANSGA region, as H171Y mutation still abrogated the constitutive Ca^2+^ influx through hO1-ANSGA channel lacking the residues 266-301 (hO1-ANSGA-ΔCT) (**Figure 6—figure supplement 1**).

**Figure 6.**
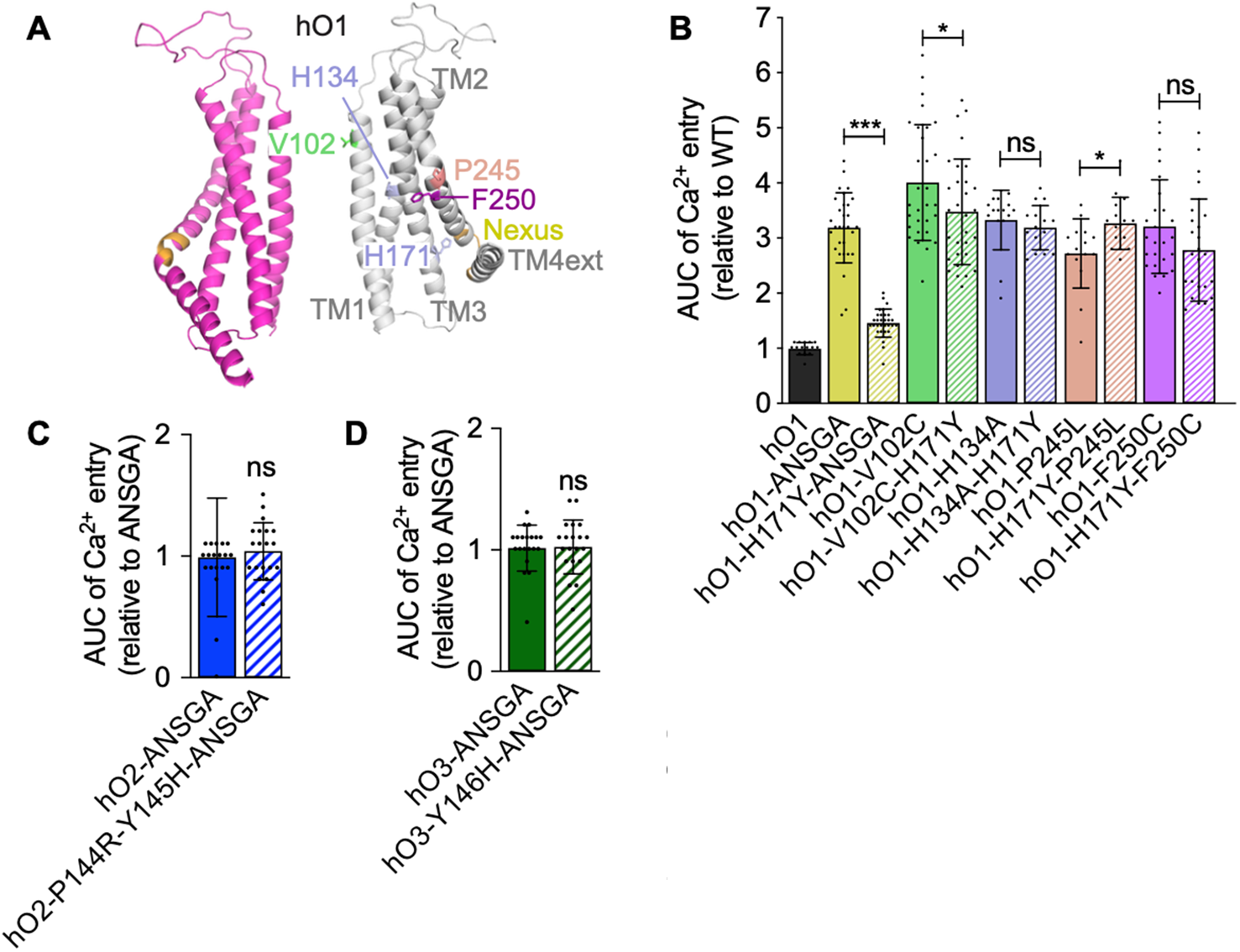
The H171Y substitution specifically inhibits the constitutive activity of hO1-ANSGA channel. **(A)** Cartoon representation of two subunits of hO1 homology model depicting studied channel-activating point mutation sites. The AUC quantified from constitutive Ca2+ entry in HEK293 cells through (**B**) different hO1 open-channel mutants with and without H171Y substitution (**C**) hO2-ANSGA with and without P144R-Y145H substitution and (**D**) hO3-ANSGA with and without Y146H substitution (mean ± SD; n ≥ 15). *p* ≤ 0.001 is indicated as “***”, 0.01 < *p* < 0.05 as “*” and *p* ≥ 0.05 as “ns”.

Finally, we addressed whether reverting the native tyrosine in hO2- and hO3-ANSGA channels, corresponding to residue position 171 of hO1, to histidine would render these mutant channels constitutively active. We incorporated the P144R-Y145H and Y146H substitutions in the hO2- and hO3-ANSGA channels, respectively, to mimic the residues present in hO1. Interestingly, it did not restore their ability to constitutively permeate Ca^2+^ ions (**Figure 4B, Figure 6C,D**). These findings indicated, that although the identified His/Tyr substitution may contribute to the inactivity of the hO2- and hO3-ANSGA nexus mutants, there may be additional substitutions in these Orai paralogs that do not allow their nexus mutants to be constitutively active.

### Substituting TM4 residue(s) of human Orai1 by human Orai2 or Orai3 or mouse Orai1 residue(s) abolishes the constitutive activity of human Orai1-ANSGA

Unlike in case of ANSGA, the hO2 P206L and hO3 P254L mutants corresponding to hO1 P245L showed constitutive Ca^2+^ entry (**Figure 7—figure supplement 1A,B,C**). Unexpectedly, however, the hO1 F250C-analogous hO2-F211C and hO3-F259C mutants did not show constitutive activity (**Figure 7—figure supplement 1A,B,C**), leading us to believe that differences in the TM4 segment between P245 and the nexus residues may contribute to the different outcomes of the ANSGA mutants of human Orai paralogs (**Figure 7A**). To address this, we first replaced several non-conserved residues spanning 246-255 positions within hO1-ANSGA by the residues present in either hO2 or hO3. The resulting hO1-F246V-I251V-A254T-V255I-ANSGA (mimicking hO2-ANSGA) and hO1-F246V-I249V-I251V-V252A-V255L-ANSGA (mimicking hO3-ANSGA) showed drastically lower or no constitutive Ca^2+^ influx in comparison to the hO1-ANSGA channel (**Figure 7B,C**). In the following experiments, we started systematically re-introducing these substitutions into a hO1-ANSGA background to pinpoint the role of individual amino acids in preventing the ANSGA-mediated conformational transition leading to hO1 channel opening. First, we limited the introduced substitutions to F246V and I251V, mimicking both hO2 and hO3, since these are the only two common residues in hO2 and hO3 within the TM4 segment corresponding to residues 246-255 of hO1 (**Figure 7A**). The I251V substitution alone did not affect the constitutive activity of the hO1-ANSGA channel (**Figure 7D**), whereas the F246V substitution completely prevented its constitutive activation (**Figure 7E,F**). Consistently, the hO1-F246V-ANSGA mutant showed a major drop in percentage of cells showing nuclear NFAT1 translocation compared to the hO1-ANSGA channel (**Figure 7G, Figure 7—figure supplement 1D**). Importantly, the F246V mutation did not affect the surface expression of the hO1-ANSGA protein (**Figure 7—figure supplement 1G**). Interestingly, mO1 contained a cysteine at the residue position 249 corresponding to F246 in hO1 (**Figure 7A**), and F246C substitution in hO1-ANSGA also abolished its constitutive activity but not the surface expression (**Figure 7H, Figure 7—figure supplement 1G**). On the other hand, the constitutive activity of the more conservative variant hO1-F246Y-ANSGA was preserved (**Figure 7H**). Also, a consistent effect of F246C and F246Y substitutions in hO1-ANSGA was observed in the NFAT1 nuclear translocation experiments (**Figure 7I, Figure 7—figure supplement 1D**). Based on these findings we hypothesized that the substitution of C249 residue at the position of hO1 F246 (**Figure 7A**). Indeed, mO1-ANSGA was not constitutively active (**Figure 7J**). However, additional C249F substitution resulted in the activation of the mO1-C249F-ANSGA but not the mO1-C249F channel, consistent with our hypothesis (**Figure 7J**).

**Figure 7.**
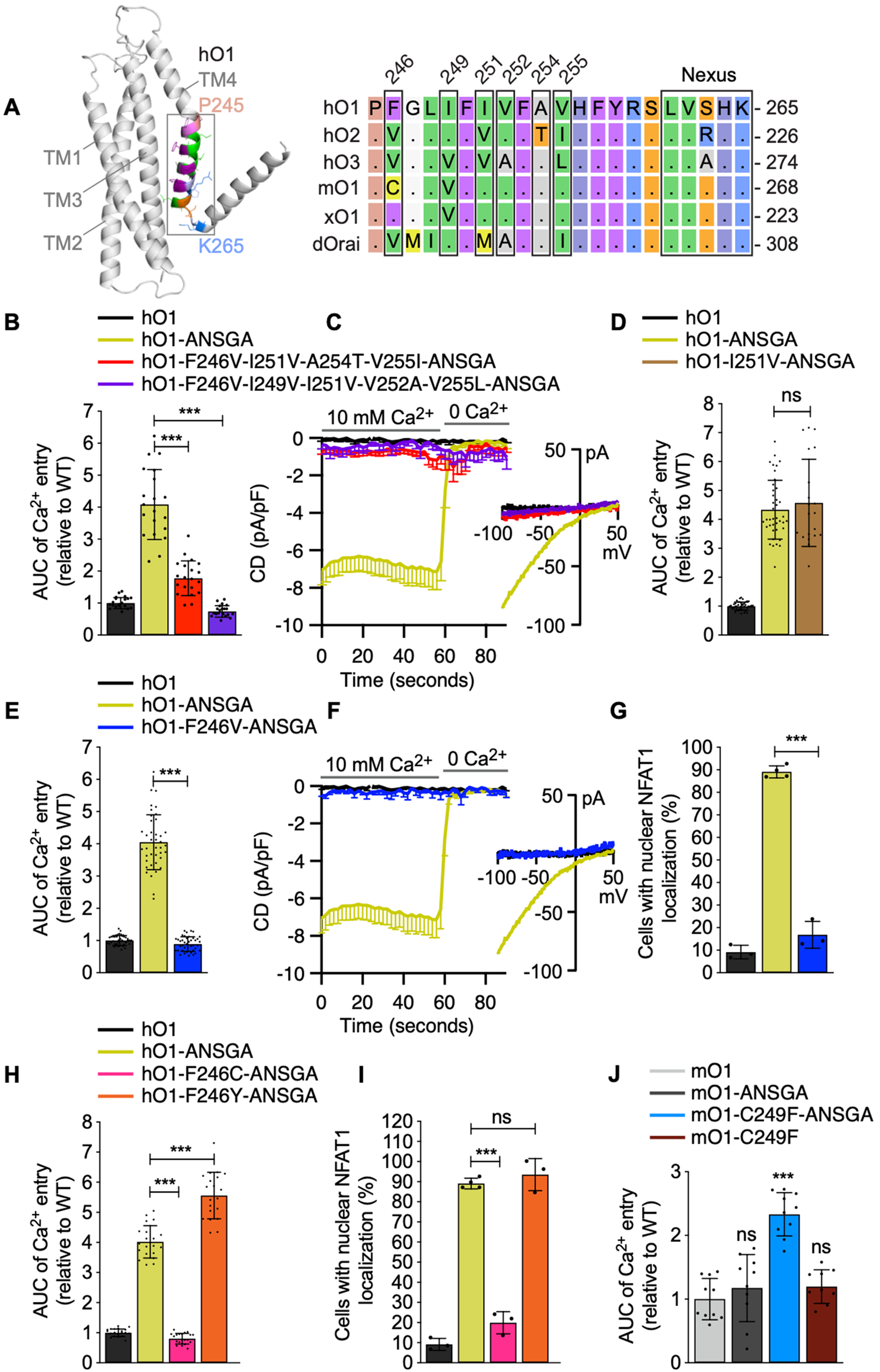
Identification of F246 residue as a gating checkpoint controlling the constitutive activity of the hO1-ANSGA channel. (**A**) Cartoon representation of a single subunit of hO1 model depicting TM4 residues P245-K265 along with the multiple sequence alignment of mentioned residues of hO1, hO2, hO3, mO1, xO1 and dOrai is shown. The highlighted residues in boxes are numbered on top according to corresponding hO1 positions. (**B**) The quantified AUC of constitutive Ca2+ entry recorded in HEK293 S1/S2 DKO cells expressing WT hO1, hO1-ANSGA and indicated hO2-mimicking and hO3-mimicking TM4 segment hO1-ANSGA mutants (mean ± SD; n=20). *p* ≤ 0.001 is indicated as “***”. (**C**) Current densities (CD) of the constitutive Ca2+ currents recorded from HEK293 S1/S2 DKO cells transiently overexpressing: WT hO1 (n=6), hO1-ANSGA (n=23), hO1-F246V-I251V-A254T-V255I-ANSGA (n=8) and hO1-F246V-I249V-I251V-V252A-V255L-ANSGA (n=8), presented as average values, -SEM, with corresponding average current-voltage (I/V) relations extracted at t= 59s. (**D**) The quantified AUC of constitutive Ca2+ entry recorded in HEK293 S1/S2 DKO cells expressing WT hO1, hO1-ANSGA and hO1-I251V-ANSGA mutants (mean ± SD; n ≥ 20). *p* ≥ 0.05 indicated as “ns”. (**E**) The AUC quantified from constitutive Ca2+ entry in HEK293 S1/S2 DKO cells through hO1-ANSGA and hO1-F246V-ANSGA channel (mean ± SD; n ≥ 20). *p* ≤ 0.001 is indicated as “***”. (**F**) Current densities (CD) of the constitutive Ca2+ currents recorded from HEK293 S1/S2 DKO cells transiently overexpressing: WT hO1 (n=6), hO1-ANSGA (n=23) and hO1-F246V-ANSGA (n=6), presented as average values, -SEM, with corresponding average current-voltage relations extracted at t= 59s. (**G**) HEK293 cells expressing WT hO1, hO1-ANSGA and hO1-F246V-ANSGA constructs having NFAT1-CFP localized to the nucleus shown as percentage (mean ±SD; n ≥ 3). *p* ≤ 0.001 is indicated as “***”. (**H**) The quantified AUC of constitutive Ca2+ entry recordings from HEK293 S1/S2 DKO cells expressing hO1-ANSGA, hO1-F246C-ANSGA and hO1-F246Y-ANSGA mutants relative to WT hO1 (mean ± SD; n=20). *p* ≤ 0.001 is indicated as “***”. (**I**) HEK293 cells expressing WT hO1, hO1-ANSGA, hO1-F246C-ANSGA and hO1-F246Y-ANSGA with nuclear localization of NFAT1-CFP are shown as percentage (mean ±SD; n ≥ 3). *p* ≤ 0.001 is indicated as “***” and *p* ≥ 0.05 as “ns”. (**J**) The AUC of constitutive Ca2+ influx recorded in HEK293 S1/S2 DKO cells expressing WT mO1, mO1-ANSGA, mO1-C249F-ANSGA and mO1-C249F mutants (mean ± SD; n = 10). *p* ≤ 0.001 is indicated as “***” and *p* ≥ 0.05 as “ns”.

We next systematically evaluated the impact of other hO2- or hO3-like substitutions within the hO1 249-255 residues segment on the constitutive activity of hO1-ANSGA nexus mutant. The combined A254T and V255I mutagenesis to mimic hO2-like residues in hO1-ANSGA did not abolish the constitutive Ca^2+^ influx through the resulting hO1-A254T-V255I-ANSGA mutant channel (**Figure 7—figure supplement 1E,F)**. On the contrary, the combination of I249V, V252A and V255L mutations to mimic hO3-like residues in hO1-ANSGA abolished the constitutive activity of hO1-I249V-V252A-V255L-ANSGA mutant channel but not its cell surface expression (**Figure 7—figure supplement 1E,F,G)**. Individual I249V, V252A, V255L or combined I249V-V255L substitutions caused only minor decrease in the constitutive Ca^2+^ influx through the hO1-ANSGA channel indicating an additive effect of these mutations in abolishment of the constitutive activity of hO1-ANSGA (**Figure 7—figure supplement 2A**). Consistently, the hO1-ANSGA mutants with individual I249V, V252A, V255L or combined I249V-V255L substitutions showed a smaller drop in percentage of cells with nuclear translocation of NFAT1 compared to the hO1-I249V-V252A-V255L-ANSGA mutant channel (**Figure 7—figure supplement 2B,C**). Altogether, these findings highlight the relevance of TM4 residues in the constitutive activation of the hO1-ANSGA channel.

### The F246V/C substitution in human Orai1-ANSGA affects the coupling between TM4 and TM3 as well as **α-**helical propensity of the TM4 helix extension

Next, to understand the mechanism of F246V-mediated inhibition of the hO1-ANSGA channel, we carried out a set of MD simulations to determine whether the F246 substitutions affect the helicity of the TM4 helix extension spanning residues 259-269. The F246V mutation in hO1-ANSGA caused a reduction in the α-helical propensity of the residues 260-264 which is also evident for the F246C mutation except for residue 262 (**Figure 8B**). Contrarily, the α-helicity of hO1-F246Y-ANSGA showed a similar trend as hO1-ANSGA. Continuing the MD analysis further, we investigated whether the F246 substitutions affect the TM4-TM3 coupling within the hO1-ANSGA nexus mutant channel (**Figure 8A**). While the mutation F246V in the hO1-ANSGA channel decreased the contact frequency between A175 and Y258 only slightly, F246C showed considerably lower A175-Y258 contacts. On the other hand, the F246Y substitution in hO1-ANSGA did not affect the contacts between A175 and Y258 (**Figure 8C**). The L174-L/A261 or F178-F257 contacts remained unchanged upon any of the F246V/C/Y mutations in hO1-ANSGA (**Figure 8—figure supplement 1**). In summary, our simulations suggest that substitution of F246 in TM4 of the hO1-ANSGA channel to valine or cysteine, but not tyrosine, affects the α-helicity of the TM4 helix extension spanning residues 259-269 as well as disrupts the local TM3-TM4 coupling.

**Figure 8.**
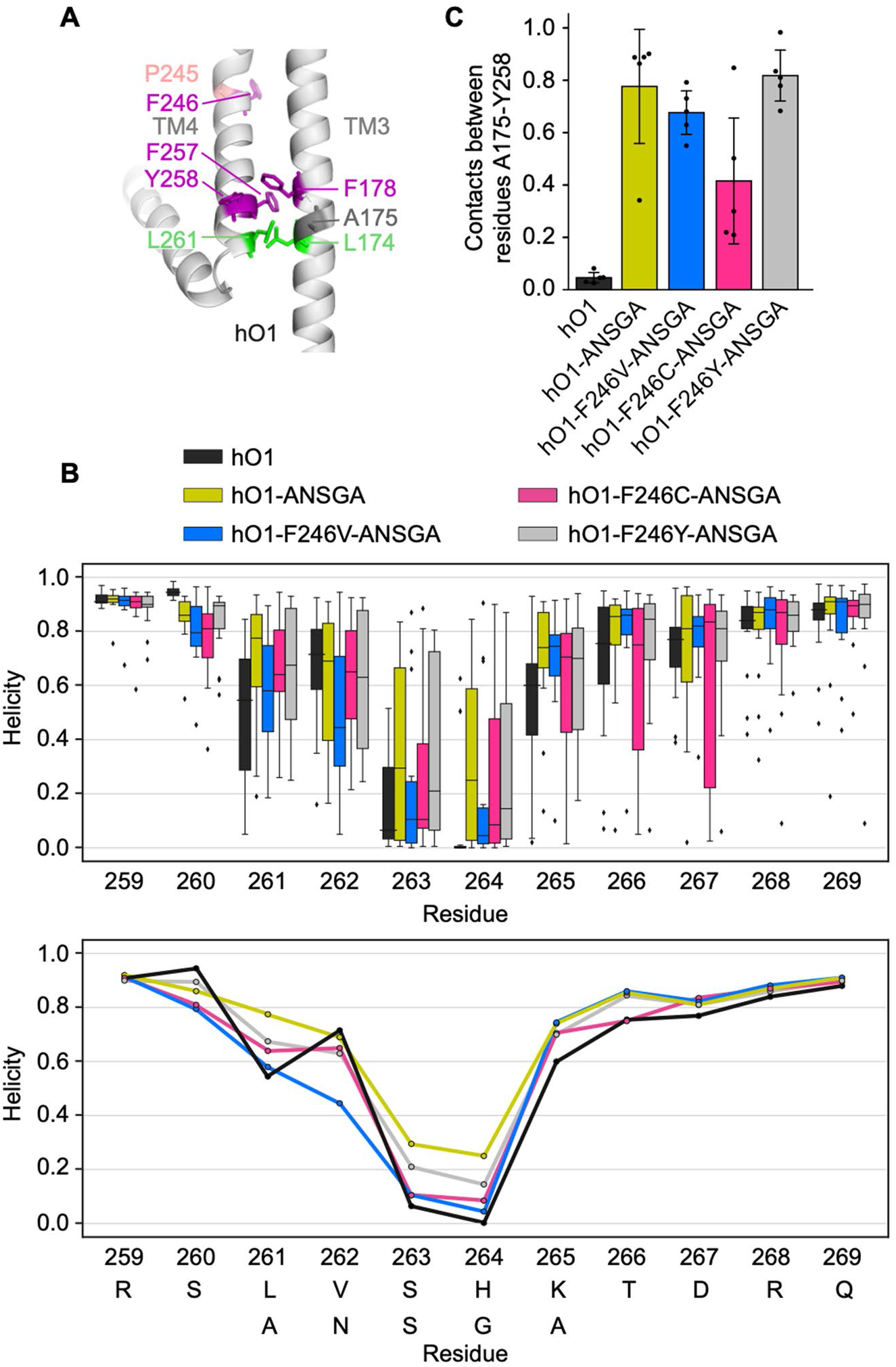
Effects of substitutions at residue position 246 on TM3-TM4 contacts and TM4-TM4ext helicity of the ANSGA channel. (**A**) Structural model showing the relevant residues in the TM3-TM4 region of hO1. (**B**) Helicity of the nexus region averaged over 5 MD trajectories for chains A, C and E, giving 15 points per residue and system (see Methods). In the upper panel, box-and-whiskers plot showing quartiles is used, outliers are plotted individually. In the lower panel, the same data is plotted for clarity using mean values. (**C**) Intra-subunit contact frequencies between residues A175 and Y258 averaged over each of the 5 MD trajectories.

### The F246V substitution does not abolish the constitutive activity of human Orai1 V102C, H134A and P245L mutants, but abolishes the constitutive activity of human Orai1-F250C mutant

In order to investigate whether the abrogating effect of the F246V substitution is specific to the ANSGA-dependent constitutive activity of the hO1, we introduced the F246V mutation into other constitutively active variants of hO1, namely the V102C, H134A, P245L and F250C mutants, as we have done earlier for H171Y (**Figure 9A**). Unlike H171Y, the F246V substitution significantly diminished the constitutive Ca^2+^ influx through the above-mentioned mutants with hO1-F246V-F250C showing no constitutive activity (**Figure 9B**). Similar to H171Y, the F246V mutation completely blocked the constitutive activity of the hO1-ANSGA-ΔCT mutant (**Figure 9—figure supplement 1**) indicating the dispensability of residues 266-301 of hO1-ANSGA in the F246V-mediated block of constitutive activity.

**Figure 9.**
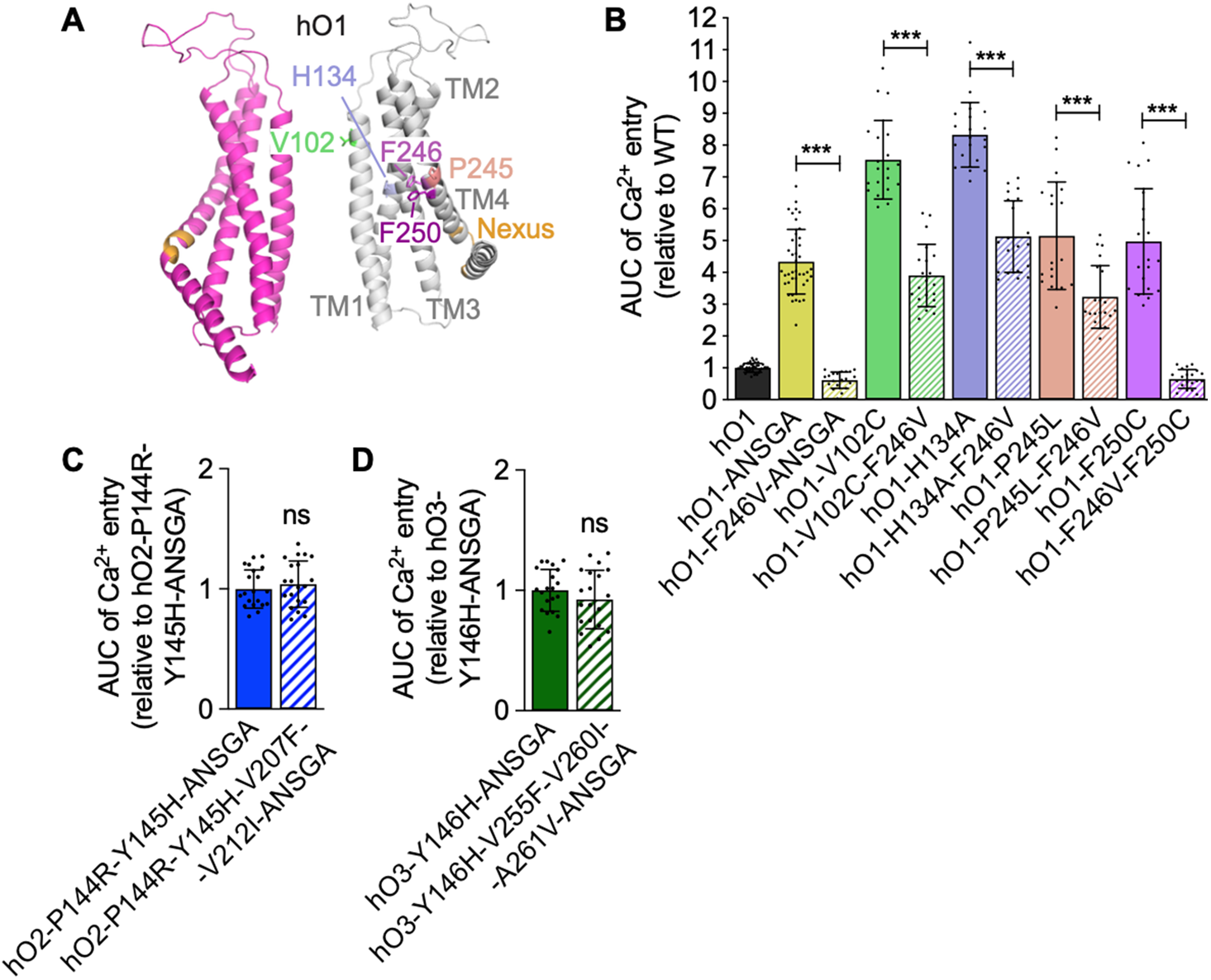
Impact of F246V mutation on the constitutive activity of ANSGA, V102C, H134A, P245L and F250C mutants of hO1. **(A)** Cartoon representation of two subunits of hO1 homology model depicting V102, H134, P245, F246 and F250 residues. Nexus region is also indicated. (**B**) The AUC quantified from constitutive Ca2+ entry in HEK293 S1/S2 DKO cells through V102C, H134A, P245L, F250C and ANSGA hO1 mutants, without and with additional F246V substitution (mean ± SD; n=20). AUC quantifications of constitutive Ca2+ measurements in HEK293 S1/S2 DKO cells expressing (**C**) hO2-P144R-Y145H-V207F-V212I-ANSGA or (**D**) hO3-Y146H-V255F-V260I-A261V-ANSGA mutants (mean ± SD; n=20). *p* ≤ 0.001 is indicated as “***” and *p* ≥ 0.05 as “ns”.

As we have identified the F246V (hO2- and hO3-mimicking) and the I249V-V252A-V255L (hO3-mimicking) substitutions in hO1 paralogs to abrogate constitutive activity of their ANSGA variants, we were interested whether the reverse, hO1-mimicking mutations in a hO2-ANSGA and hO3-ANSGA background would enable constitutive activity of these channels. To this end, we used our hO2-P144R-Y145H-ANSGA and hO3-Y146H-ANSGA mutants as background, which already contained hO1-mimicking substitutions at positions corresponding to H171 and R170 in hO1 (**Figure 4B**). In these hO2 and hO3 constructs, we introduced further substitutions mimicking the hO1 F246 (V207F in hO2 and V255F in hO3), I251 (V212I in hO2 and V260I in hO3) and V252 (A261V in hO3) residues (**Figure 7A**). However, both of these hO1-like hO2-ANSGA and hO3-ANSGA channels remained closed (**Figure 9C,D**). In addition to the above-mentioned reversal mutations, the hO3-ANSGA mutant further containing the V258I-A261V-L264V substitutions, corresponding to the reversal of hO1 I249V-V252A-V255L mutations (**Figure 7A**) that inhibited hO1-ANSGA activity, also remained constitutively inactive (data not shown). Clearly, there are further differences among the human Orai paralogs which render hO2 and hO3 incapable of being activated by the ANSGA nexus mutation.

### Human Orai1 H171Y as well as F246V mutants can be gated by STIM1

Finally, we aimed to determine whether the H171Y and F246V mutations also interfere with the canonical STIM1-dependent activation of hO1. To this end, we generated relevant hO1 mutants without any mutations in the nexus residues, expressed them in the S1/S2 DKO HEK293 cells along with the STIM1 protein, and measured *I_CRAC_* upon stimulation with inositol triphosphate (IP_3_). Similarly, we tested the hO1-H171Y-ANSGA and hO1-F246V-ANSGA mutants. The obtained results showed that both hO1-H171Y and hO1-F246Y mutants could be activated by STIM1, while the corresponding ANSGA nexus mutants remained inactive (**Figure 10**). Notably, the hO1-H171Y mutant, gated by STIM1, displayed *I*_CRAC_ characteristics typical for the WT channel (**Figure 10A**). The hO1-F246V mutant gated by STIM1, however, evoked *I*_CRAC_ characterized by different kinetics and current size. Moreover, it exhibited a clear inactivation component, which was not present in the WT hO1- or hO1-H171Y-mediated currents (**Figure 10B**). These findings, however, require further elucidation beyond the scope of the current work.

**Figure 10.**
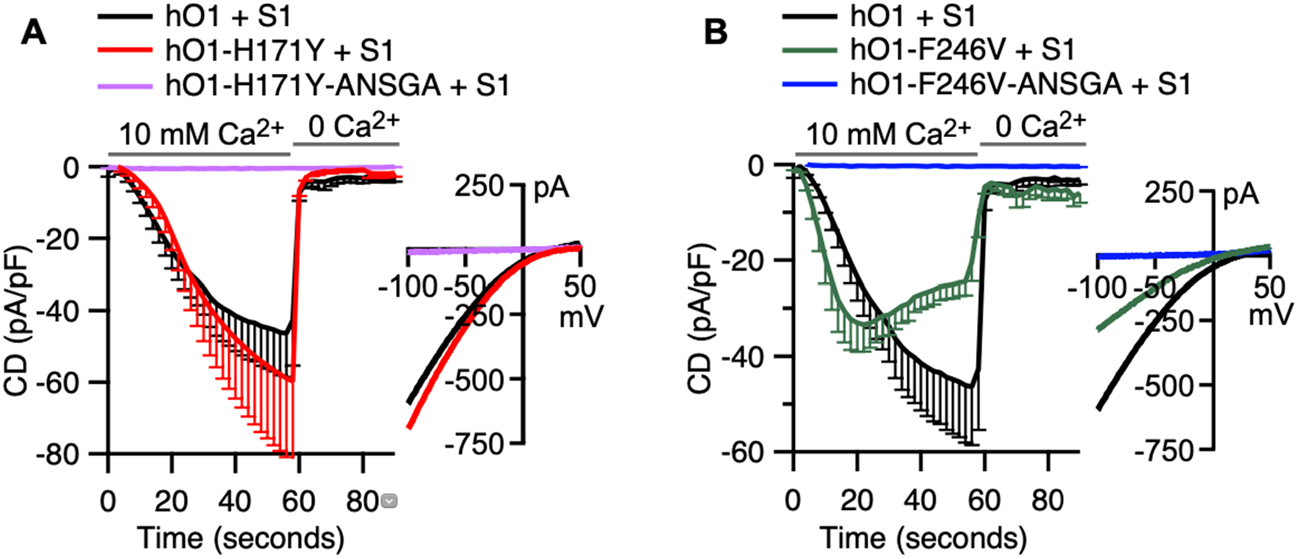
STIM1 is able to gate both H171Y and F246V mutants of hO1 as assessed by electrophysiological recordings. **(A)** Current densities (CD) of the *ICRAC* currents recorded from HEK293 S1/S2 DKO cells transiently overexpressing: WT hO1 + hS1 (n=9), hO1-H171Y + hS1 (n=6) and hO1-H171Y-ANSGA + hS1 (n=6), presented as average values, -SEM, with corresponding average current-voltage (I/V) relations extracted at t= 59s. (**B**) Current densities (CD) of the *ICRAC* currents recorded from HEK293 S1/S2 DKO cells transiently overexpressing: WT hO1 + hS1 (n=9), hO1-F246V + hS1 (n=7) and hO1-F246V-ANSGA + hS1 (n=6), presented as average values, -SEM, with corresponding average current-voltage (I/V) relations extracted at t= 59s.

## Discussion

In recent years, several studies have put forward different models for hO1 channel activation, which include the rotation of the TM1 helix (Yamashita et al., 2017), twist-to-open (Dong et al., 2019), counter-ion assisted permeation (Liu et al., 2019) and a pore dilation model (Frischauf et al., 2017; Hou et al., 2018; Hou et al., 2020). However, the exact gating mechanism of the hO1 channel by STIM1 is far from being fully understood and is awaiting structural studies of the STIM1-Orai1 complex. Nevertheless, numerous studies have led to the current understanding that the binding of STIM1 at the C-terminus of Orai1 initiates a series of conformational changes that propagate through numerous gating checkpoints within Orai1 subunits and thereby culminate in CRAC channel activation (Fahrner et al., 2018; Frischauf et al., 2017; Palty et al., 2015; Tiffner et al., 2020; Yeung et al., 2018; Y. Zhou et al., 2016). A major goal of our study was to investigate whether these checkpoints and gating elements are conserved among the three human Orai paralogs.

We first investigated whether the TM1-TM2 conformational coupling was also conserved in hO2 and hO3 by mutagenesis of the residues corresponding to H134, a “checkpoint” residue of this coupling in hO1 (Frischauf et al., 2017; Yeung et al., 2018). The constitutive Ca^2+^ current observed in both hO2-H108A and hO3-H109A (**Figure 2C**) suggests a conserved TM1-TM2 interplay in the gating of human Orai channels. Further, constitutive Ca^2+^ influx through dOrai-H206A was reported recently (Hou et al., 2020). Moreover, in line with our data, a recent report published during the preparation of our manuscript demonstrated constitutive current through the H109A mutant of hO3 but did not test the same for hO2 (Tiffner et al., 2021). Also, the study of Tiffner *et al*. did not investigate if hO3-H109A or hO2-H108A would lead to nuclear translocation of NFAT1 (**Figure 2D,E**). On the other hand, the H134A mutant of hO1 has been shown to effectively cause the translocation of NFAT1 to the nucleus (Frischauf et al., 2017). In summary, our results point in the direction that the sequence of events during the gating mechanism downstream of the H134 checkpoint are conserved in a wide range of Orai proteins.

While both earlier studies related to the H134 checkpoint suggest a conformational coupling mechanism between TM2 and TM1 of hO1 leading to channel opening, different mechanistic explanations have been proposed. Frischauf *et al*. suggest that TM2 H134 forms a hydrogen bond with the TM1 S93 residue and this serves as a hydrogen bond-dependent trigger that gates hO1 channels (Frischauf et al., 2017). On the contrary, Yeung *et al*. argue against the formation of this hydrogen bond and rather suggest that substitution of the bulky side chain of H134 by smaller amino acids allows rotation of the TM1 F99 away from the pore axis (Yeung et al., 2018), a phenomenon that was not observed by Frischauf and co-workers (Frischauf et al., 2017). Although the rotation of the TM1 helix was also not evident from the recent structure of dOrai-H206A (corresponding to hO1-H134A), the rigid-body outward movement of each dOrai-H206A subunit resulted in repositioning of F171 (corresponding to hO1 F99), which slides away from the pore, widening the hydrophobic region (Hou et al., 2020). In contrast, we could observe the TM1 (F99) rotation in hO1-H134A MD simulations as well as reproduce our earlier findings using a homology-based model of hO1 bearing the H134S substitution (Bulla et al., 2019) (**Figure 5—figure supplement 2A**). Overall, our MD simulations confirm that sustained Orai1 channel opening requires the displacement of F99 away from the pore.

Our results also indicate that the mechanistic function of the sulfur-aromatic latch is conserved in all three human Orai channels. The M101 residue in hO1 TM1 has been suggested to function as a “gate latch” by engaging the hydrophobic gate (F99) of the neighboring subunit to sustain the open conformation of the hexameric channel (Yeung, Ing, et al., 2020). Consistently, our MD simulations have confirmed a higher contact frequency between F99-M101 in hO1-H134A/S compared to the WT channel (**Figure 2—figure supplement 2C,D,E**). Our finding is in line with the report on narrowing of the distance between F171 and M173 of dOrai (corresponding to hO1 F99 and M101, respectively) in the open state of the channel (dOrai H206C/Q) compared to WT (Yeung, Ing, et al., 2020). In the other, resting state of the “gate latch”, it has been suggested that M101 partners with F187 in the TM3 of the adjacent subunit, stabilizing the M101-F187 resting interaction. This, in turn, may arrest the hO1 channel in a closed state, likely due to the release of F99 side chain into the pore-facing conformation (Yeung, Ing, et al., 2020). Interestingly, hO1-F187C was reported constitutively active (Yeung et al., 2018) and in light of the recent findings from Prakriya’s laboratory, it is reasonable to envision that due to the inability of C187 to engage in the M101-dependent gate latch, M101 instead permanently arrests F99 out of the pore axis, supporting the constitutively active state of the hO1-F187C channel (Yeung, Ing, et al., 2020). Our findings that the hO1-F187C and the corresponding hO2-F161C and hO3-F162C channels are constitutively active (**Figure 2—figure supplement 2F,G,H,I**) suggest that the recently proposed M101-F187 interactions which support hO1 channel closing (Bonhenry et al., 2021; Yeung, Ing, et al., 2020) are also conserved in hO2 and hO3. Overall, our data show that the inter-subunit conformational coupling of hO1 between TM1-TM2 (F99-H134), as well as TM1-TM3 (M101-F187), are also conserved in hO2 and hO3 paralogs.

Moving more distal from the pore, we showed that the TM4 mutations corresponding to the position hO1 P245, namely P206C and P254C in hO2 and hO3, respectively, rendered them constitutively active (**Figure 7—figure supplement 1A,B,C**). Consistently, constitutive current through hO3-P254L has also been recorded in a previous report (Derler et al., 2018). The structure of P288L dOrai (corresponding to P245L hO1) shows a clear straightening of the kink in TM4 at P288 (Liu et al., 2019). The conserved constitutive activation effect of P245, P206 and P254 substitutions in hO1, hO2 and hO3, respectively, suggests that these mutations may trigger similar structural rearrangements among Orai paralogs. In our MD simulations, the tubular aggregate myopathy (TAM)-associated P245L mutant showed a significantly higher α-helical propensity of the TM4 fragment upstream of L245 (residues 241-244) compared to WT (**Figure 5—figure supplement 1A**), indicating the unbending of the proline kink as a possible mechanism of how this mutation opens the channel.

In contrast to the above-mentioned gating checkpoints that we show to be conserved among human Orai channels, we also report positions that could represent steric brakes that are not conserved among the three human Orai channels. Interestingly, these positions lie further away from the pore, in the TM4 and TM4ext regions.

One of these positions is residue F250 within TM4 of hO1, the mutation of which (F250C) was identified in a cysteine screen of hO1 TMs (Yeung et al., 2018) to lead to the constitutive activation of hO1 (Tiffner et al., 2020; Yeung et al., 2018). Our data shows that unlike the mutants of hO2 and hO3 corresponding to hO1 P245L, the hO2-F211C and hO3-F259C mutants, corresponding to hO1-F250C, do not exhibit any constitutive activity (**Figure 7—figure supplement 1A,B,C**).

Importantly and central to our work, we also show that the ANSGA “nexus” mutations, further along the protein chain in the hinge region between TM4 and TM4ext, likely also exert their activating effect through a non-conserved mechanism. Based on hO1-ANSGA, we expected the ANSGA mutants of hO2 and hO3 to also display constitutive activity and NFAT1 activation. However, this was not the case (**Figure 3C,D,E,F,G,J,K**). Also, despite dOrai, xO1 and mO1 all having the LVSHK nexus residues conserved, only the ANSGA mutant of xO1 exhibited the constitutive activity (**Figure 3H,I and Figure 7J**).

Although all of the P245, F250 and the LVXHK nexus residues of hO1 are conserved among different human Orai paralogs as well as in Orai channels of lower organisms, the neighboring residues within TM4 are not very well conserved, e.g. *Drosophila* Orai1 protein, which shares only 47% overall identity with hO1, has several additional non-conserved residues downstream from P288 corresponding to P245 of hO1 (**Figure 7A**). Thus, the differences in sequence in this region might interfere with constitutive activation by ANSGA in certain Orai channels and may also explain differences in gating mechanism in various Orai paralogs and orthologs.

Current knowledge on the mechanism by which STIM1 or ANSGA nexus mutations activate Orai1 is incomplete and does not explain our observed lack of activation by the ANSGA mutation in hO2, hO3 and certain Orai orthologs. The gating-related conformational changes that initiate within the TM4 helix of hO1 upon binding of STIM1 to its C-terminus and propagate to the channel’s pore-forming TM1 presumably rely on flexion of the hO1 C-terminal nexus hinge residues (Y. Zhou et al., 2016). In support of this mechanism, we observed a notable increase in the calculated helicity of the LVSHK region when substituted with ANSGA residues, which may be indicative of the flexion of TM4ext (**Figure 5B**). Disrupting the flexibility of the LVSHK hinge, for example by the S263P mutation, abolished *I*_CRAC_ (Tirado-Lee et al., 2015) even when hO1 was covalently linked with two active fragments of STIM1, suggesting a critical role of hinge flexibility in gating of hO1 by STIM1 (Palty et al., 2015). The motifs corresponding to LVSHK in hO2 and hO3 contain either an arginine or an alanine residue, respectively, in the place of serine (**Figure 3B**). These substitutions, however, are not expected to change the overall structure or flexibility of the LVSHK hinge, as the mutation of S263 to either arginine or alanine did not affect the interaction between hO1 and channel activation domain of STIM1 (Tirado-Lee et al., 2015). Therefore, hO2 and hO3 are envisioned to have an overall similar structural architecture of the nexus hinge as hO1. Furthermore, mutations of hO2 and hO3 residues corresponding to the earlier mentioned L273 and L276 residues of hO1 also affected the interaction of hO2 and hO3 with STIM1, as well as diminished the *I*_CRAC_ (Baraniak et al., 2021; Frischauf et al., 2009; Frischauf et al., 2011; Niu et al., 2020). The mechanism how ANSGA causes the constitutive activity of the hO1 channel is also likely different from that of the P245L mutation, since the helicity changes observed in our MD simulations upon ANSGA substitution of LVSHK residues in hO1 remained restricted around the nexus region without affecting the helicity of the TM4 helix region (residues 240-250) encompassing the proline bend caused by P245 (**Figure 5A**, **Figure 5—figure supplement 1A**).

Interestingly, we were able to uncover three positions that seem to interfere with ANSGA-induced activation of hO1, all of which point at the importance of TM4-TM3 contacts in transducing gating-relevant conformational changes from the distal TM rings towards the central pore. The first of these positions, which we have identified through multiple sequence alignments and homology-based modeling, is H171, located in the cytosolic extension of TM3 helix (TM3ext), proximal to the LVSHK motif, and not conserved in hO2, hO3 and dOrai (**Figure 4A,B**). When H171 was substituted by tyrosine, which is the native residue at the corresponding position in hO2, hO3 and dOrai, both the constitutive activation of hO1-ANSGA and NFAT1 activation were lost (**Figure 4C,D,G**). Surprisingly, however, the H171Y substitution did not eliminate the constitutive activity of other known hO1 open channel mutants such as TM1 V102C (McNally et al., 2012; Yamashita et al., 2017), TM2 H134A (Frischauf et al., 2017; Yeung et al., 2018), TM4 P245L (Nesin et al., 2014; Palty et al., 2015) and TM4 F250C (Yeung et al., 2018) (**Figure 6A,B**). Similarly, the H171Y substitution in hO1 with native nexus “LVSHK” residues did not affect STIM1-mediated gating of the channel (**Figure 10A**). Nevertheless, the importance of the H171 residue in gating is also corroborated by a recent report showing that the H171D substitution, identified through randomized mutations, minimized the activity of the engineered light-operated hO1-P245T channel in dark, while retaining its activity under blue light illumination (He et al., 2021). Residue H171 of hO1 appears to be a feature in the Orai1 proteins of vertebrates, whereas a conserved tyrosine residue takes the corresponding position in invertebrates. A study suggesting diversification of Orai gating mechanisms between invertebrates and mammals showed that the F192R-Y193H substitution in the Orai channel of *C. elegans* to mimic hO1 residues R170-H171 led to STIM1-independent constitutive activation of *C. elegans* Orai channel, as well as nuclear translocation of NFAT1 (Kim et al., 2018). Thus, we propose that the H171 position is essential to Orai channel gating, nevertheless its role in hO1 might be limited to local, “weak” stabilization of TM3-TM4 contacts. Thus, a H171Y substitution, native to hO2 and hO3 at this position, abrogates the effect of the ANSGA mutation in hO1, but not that of other activating mutations, and it also does not affect normal STIM-mediated gating.

In pursuit of understanding the mechanism of action of the H171Y-mediated selective inhibition of hO1-ANSGA constitutive activity, we show that TM3-TM4 contacts are essential for ANSGA-induced activation. In particular, TM3-TM4 interaction pairs F178-F257 (Liu et al., 2019) and L174-L/A261 (Y. Zhou et al., 2016) have been previously shown to be important for hO1 channel activation, and we found that these residues, as well as Y258 (TM4) are indispensable for the activity of the hO1-ANSGA channel as well (**Figure 5D,E and Figure 5—figure supplement 1C**). While the ANSGA substitution did not affect the contact frequencies of these TM3-TM4 residue pairs in our MD simulations, it did result in significantly enhanced contact frequencies of another TM3-TM4 residue pair, A175 and Y258, which were in turn impaired by a further H171Y substitution (**Figure 5A,C**). Furthermore, we found that all the residues past the ANSGA sequence do not take part in shaping the constitutive activity of the channel, since the constitutive activity of hO1-ANSGA-ΔCT channel was not compromised (**Figure 6—figure supplement 1 and Figure 9—figure supplement 1**), which is consistent with earlier reports (L. Zhou et al., 2018; Y. Zhou et al., 2016). In addition, the H171Y mutation still inhibited the constitutive activity of the hO1-ANSGA channel even after the removal of residues 266-301 (**Figure 6—figure supplement 1**). Therefore, even though the H171Y substitution reduced the α-helical propensity of hO1-ANSGA residues 265-267 C-terminal to the ANSGA motif (**Figure 5B**) in our MD simulations, the last 36 residues beyond residue 265 are dispensable for ANSGA-induced gating. Based on these results, we believe that the impairment in the local coupling between TM3-TM4 (A175-Y258) may be the relevant reason for the disruption of hO1-ANSGA channel activity by the H171Y substitution.

The second position that we found critical in controlling the constitutive activity of hO1-ANSGA to allow influx of Ca^2+^ in a constitutive manner with subsequent nuclear translocation of NFAT1 was residue F246. F246 in hO1 is conserved in xO1 but neither in hO2, hO3 and dOrai, where it is replaced by a valine residue, nor in mouse Orai1, where it is replaced by cysteine. Herein, both F246V and F246C substitutions in hO1-ANSGA prevented the constitutive Ca^2+^ influx and subsequent nuclear translocation of NFAT1 (**Figure 7E,F,G,H,I and Figure 7—figure supplement 1D**). Thus, we can speculate that the valine residue in hO2 and hO3 at the positions corresponding to F246 of hO1 participates in exerting a brake on the putative constitutive activity of the hO2- and hO3-ANSGA channels.

Although the F246V mutation did not abolish the activity of other constitutively active hO1 mutants such as V102C, H134A and P245L, it significantly decreased the measured influx of Ca^2+^. However, unexpectedly, substitution of F246 to valine completely abolished the constitutive influx of Ca^2+^ through the hO1-F250C and hO1-ANSGA mutants (**Figure 9A,B, Figure 7E**). Since F246V did not alter the surface trafficking of hO1-ANSGA (**Figure 7—figure supplement 1G**), it is likely that this mutation does not alter the trafficking of V102C, H134A, P245L or F250C mutants either. Similarly to H171Y, our MD simulations suggest that TM4 mutations (F246V/C) that interfere with the constitutive activation of hO1-ANSGA also impair local coupling between TM3-TM4 (A175-Y258) and α-helicity of the nexus segment (**Figure 8**). Since we also observed a significant decrease in the amplitude of STIM1-activated *I*_CRAC_ mediated by hO1-F246V in comparison to the WT hO1 (**Figure 10B**), it is likely that this TM4 residue is important in regulating the channel activity. In comparison to the H171 position, F246 seems to be a “strong” effector of gating in hO1, since it is able to abolish the effects of the activating mutations ANSGA and F250C, while at the same time to significantly attenuate Ca^2+^ influx mediated either by the other tested constitutively active mutant variants of hO1, or by STIM1-mediated gating.

In addition, as a third position affecting the gating of hO1-ANSGA, we found that a combination of other residue substitutions mimicking the residues of hO3 TM4 on hO1 (I249V-V252A-V255L) leads to abrogation of the constitutive activity of hO1-ANSGA (**Figure 7—figure supplement 1E,F and Figure 7—figure supplement 2**). These positions seem to have an additive effect, which implies that there could be several other positions making contributions to shape the energetic barrier of constitutive activity.

Interestingly, while the substitution of H171 by tyrosine abrogated the constitutive activity of hO1-ANSGA, the reverse substitution of the corresponding hO2 Y145 and hO3 Y146 residues by histidine was not sufficient, either alone or in combination with TM4 mutations, to evoke a constitutive Ca^2+^ influx through hO2- or hO3-ANSGA mutant channels (**Figure 6C,D, Figure 9C,D**). These data provide further evidence that other residue differences in hO1 and hO2/hO3 can shape the overall energetics of channel gating. Mouse Orai1 protein, which shares 90% sequence identity with hO1, has a cysteine at the residue position corresponding to F246 of hO1 with most of the downstream residues conserved compared to hO1 (**Figure 7A**). Strikingly, the ANSGA mutant of mO1, which was otherwise constitutively inactive, resulted in a modest constitutive activation when C249 was substituted with a phenylalanine residue to mimic the native residue (F246) of hO1 (**Figure 7J**). This interesting result suggests that at least in certain Orai channels, the energetic barrier of activation can be lowered enough by a single point mutation so that activating mutations such as ANSGA can induce constitutive activity in the channel.

Earlier, it was shown that the addition of the ANSGA substitution in the otherwise non-selective hO1-V102C pore mutant channel renders selectivity towards Ca^2+^ and that the mutations that abrogate STIM1-mediated activation of hO1, such as the L174D mutation, also eliminate the constitutive activity of hO1-ANSGA channel (Y. Zhou et al., 2016). Based on these findings it was suggested that the ANSGA substitution of the “LVSHK” nexus residues of hO1 mimics the STIM1 gating trigger and causes a conformational change in the hO1 TM4ext leading to the opening of the channel pore formed by TM1. However, we show here that two different hO1 single point mutations, one in TM3ext (H171Y) and another in TM4 (F246V), adopted from other Orai homologs, completely abolish the constitutive activity of the O1-ANSGA channel but do not shut down the permeation of Ca^2+^ through STIM1-gated hO1 channels. This provokes us to reconsider how well the ANSGA mutant resembles the actual STIM1-gated open state of hO1. As the conformational trigger by the ANSGA substitution leading to the opening of hO1 channel is obstructed by changes that still allow gating by STIM1, it is likely that either the ANSGA channel does not follow the same order of conformational rearrangements that lead to the channel opening, or that the gating stimulus provided by STIM1 supersedes the stimulus provided by ANSGA (**Figure 11)**. In other words, we could expect the ANSGA nexus mutations to be “weak” activators, while STIM1 binding provides a “strong” stimulus that is able to overcome higher energetic barriers to gate the channel as depicted in our hypothetical model (**Figure 11B,C**). We propose that mutations of TM3ext H171 and TM4 F246 in hO1 interfere with the ANSGA-mediated gating signal by weakening the local coupling between TM3 and TM4 (**Figure 11E**). Nevertheless, the stimulus exerted by STIM1 binding is strong enough to open the H171Y and F246V mutant hO1 channels with intact nexus residues, regardless of the weakened TM3/TM4 coupling (**Figure 11D**). While our computational efforts also provide mechanistic insights into how the relevant H171 and F246 substitutions exert a “brake” on the relay of conformational changes along the gating pathway, future experimental studies are needed to further elucidate their exact mechanism of action.

**Figure 11.**
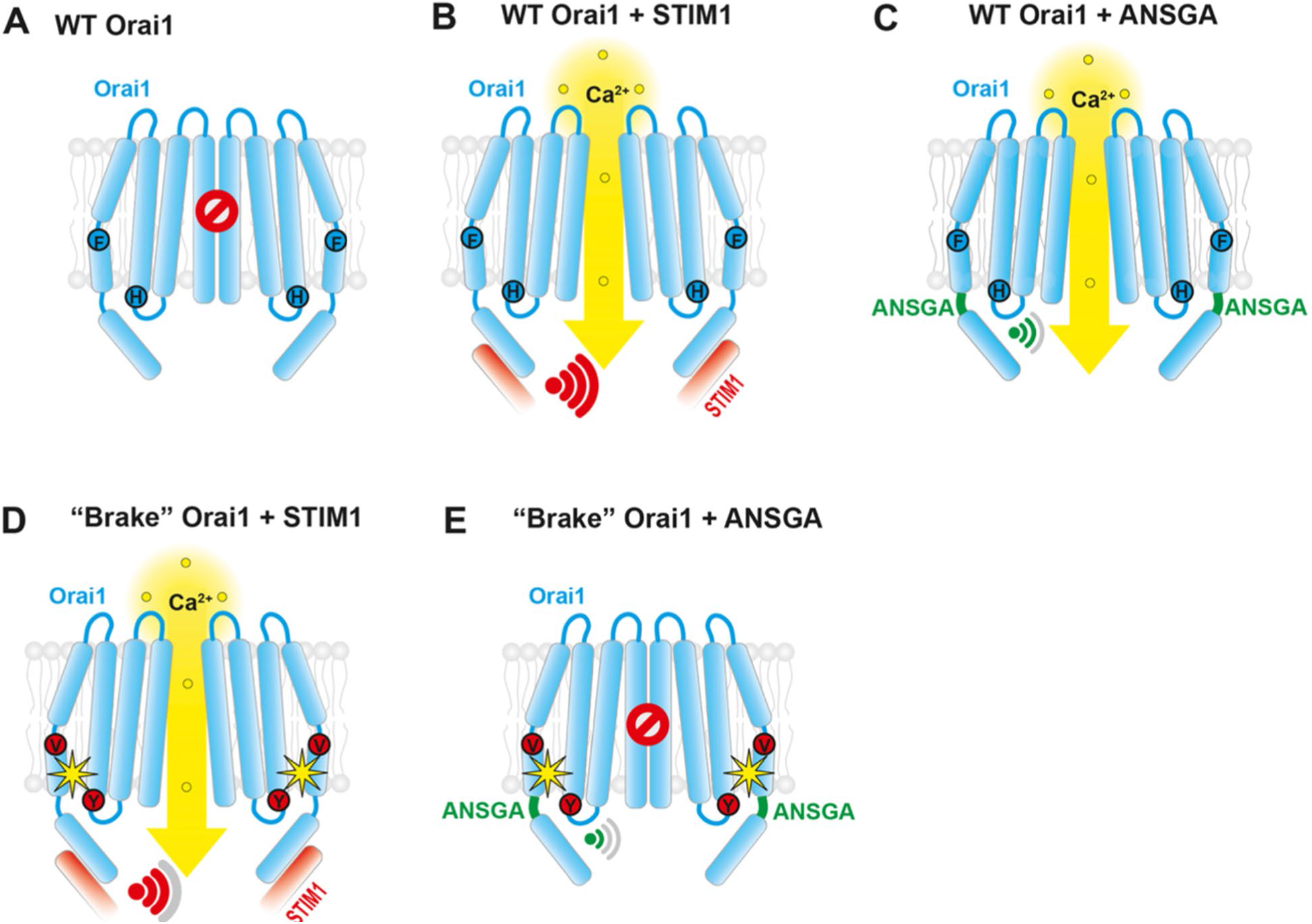
Hypothetical mechanism of hO1-channel activation by STIM1 and the “ANSGA” nexus mutation. (**A**) hO1 in a closed, quiescent state without any gating stimulus. (**B**) STIM1 binds the C-terminal extension (TM4ext) of hO1, and exerts a strong gating signal that propagates through TM3-TM4 contacts to open the central pore. (**C**) The ANSGA mutation exerts a moderate gating signal that is able to open the hO1 channel through intact TM3-TM4 contacts. (**D**) Brake mutations at positions H171 and F246 introduced in hO1 interfere with the gating signal by weakening the TM3-TM4 contacts, nevertheless, the stimulus exerted by STIM1 binding is still strong enough to open the channel. (**E**) Brake mutations attenuate the gating signal by ANSGA sufficiently enough to hinder the opening of the Orai channel.

We envision that our results on the various Orai channel isoforms could serve as a source of information for structural biology and structure-based drug design. Structural insights into Orai1 channel activation have been generated using the *Drosophila* TM4 mutant P288L corresponding to hO1-P245L and the *Drosophila* TM2 mutant H206A corresponding to hO1-H134A (Hou et al., 2018; Hou et al., 2020; Liu et al., 2019). Our study indicates that mouse O1 ANSGA and *Xenopus* O1 ANSGA channels exhibit “intermediate-arrested” or “fully-activated” states that could be used for future structural work to gain further insights into hO1 channel activation and pharmacological inhibition. Several potent inhibitors of human Orai channels have been reported in the literature (Bakowski, Murray, & Parekh, 2021; Schild et al., 2020), but none of them exhibits remarkable Orai-subtype specificity. Our work could help design strategies to generate Orai structures in complex with inhibitors, as a basis of structure-based design of Orai subtype-specific channel modulators.

## Materials and Methods

### Cloning and site-directed mutagenesis

*Xenopus* Orai1 cDNA (Clone ID: 4633914, MXL1736-202771770, Horizon Discovery Ltd.) after PCR amplification was subcloned into pmCherry-C1. Human Orai2 was amplified by PCR from CFP-Orai2 peCFP-C1 provided by Prof. Christoph Romanin, University of Linz, Austria and sub-cloned into pmCherry-C1. Mouse Orai1 WT was constructed by site-directed mutagenesis-mediated generation of a stop codon after the coding sequence of Orai1 in mouse WT Orai1-eGFP peGFP-N1 provided by Prof. Francisco Javier Martin-Romero, University of Extremadura, Spain. All the forward primers used for site-directed mutagenesis are mentioned in the key resources table and the reverse primers used were reverse complementary. Pfu Ultra DNA polymerase (600380, Agilent Technologies) was used for site-directed mutagenesis.

### CRISPR/Cas9-mediated generation and validation of STIM1/STIM2 double knockout HEK293 cells

To generate STIM1/STIM2 double knockout HEK293 cells using CRISPR/Cas9 genome editing technique, we first knocked out STIM1 from HEK293 cells as described in (Butorac et al., 2019). Briefly, two different guide RNA (gRNA) were designed using the Benchling CRISPR webtool (https://benchling.com/crispr) that target exon 1 of STIM1. The gRNA1 (5’-TTC TGT GCC CGC GGA GAC TC-3’) was chosen to target a region encoding 5’ untranslated region (UTR) of STIM1 and gRNA2 (5′-GTA TGC GTC CGT CTT GCC CTG-3′) to target a region encoding the signal peptide of hSTIM1. Each of the gRNAs were cloned into pX330.pgkpuro vector (Harmsen, Klaasen, van de Vrugt, & Te Riele, 2018) and co-transfected into the HEK293 (ATCC^®^ CRL-1573™) cells. The transfected cells were selected by 96 h treatment with 1.5 µg/ml puromycin. The puromycin selected HEK293 cells were then used to knock out STIM2. STIM2 gRNA1 (5’-CGG AAC CAAT GAA CGC AGC C-3’) was chosen to target a region that is thought to encode an alternate longer signal peptide of hSTIM2 (Graham, Dziadek, & Johnstone, 2011), whereas gRNA2 (5’-GCT GGT AGC CGG AGC GGC GGA-3’) was chosen to target the regular short signal peptide encoding sequence of hSTIM2 (Bhardwaj, Hediger, & Demaurex, 2016). STIM2 gRNA1 was cloned into BbsI-HF linearized pX330.pgkpuro and gRNA2 was cloned into BbsI-HF linearized pU6-(BbsI)_CBh- Cas9-T2A-mCherry (Addgene Plasmid #64324). After co-transfection with STIM2 gRNA1 and gRNA2 plasmids the cells were sorted by Fluorescence-Activated Cell Sorting (FACS) with mCherry fluorescence and single cells were seeded in a 96-well plate.

To validate the knockout, genomic DNA was extracted from the resulting single-cell-derived clones and 5’-CCT TCC GCA GGG GTG TAG T-3’ forward and 5’-CTC CAA CAG CCA AAG GTC AA-3’ reverse primers were used to PCR amplify a 181 bp STIM1 amplicon. Similarly, 5’-CGG AAC CAA TGA ACG CAG C-3’ forward and 5’-GAG TCG AGG CGG GAT GAA G-3’ reverse primers were used to obtain a 222 bp STIM2 PCR product. Western blotting was performed as described in (Bhardwaj et al., 2020). Guinea pig polyclonal hSTIM1 antibody (Ercan, Chung, Bhardwaj, & Seedorf, 2012) was used in a 1:1,000 dilution and peroxidase-conjugated affinipure goat anti-guinea pig secondary antibody (106-035-003, Jackson Immuno Research) was used in a 1:10,000 dilution. Rabbit polyclonal hSTIM2 antibody (4917S, Cell Signaling) was used in a 1:1000 dilution and horseradish peroxidase (HRP)-conjugated goat anti-rabbit secondary IgG (W401B, Promega) was used in a 1:20,000 dilution. Mouse monoclonal Tubulin antibody (T9028, Sigma-Aldrich) was used in a 1:2,000 dilution and HRP-conjugated goat anti-mouse secondary antibody was used in a 1:3,000 dilution (172-1011, Bio-Rad). Following gDNA based PCR screening and Western blotting, SOCE recordings were performed using the S1/S2 DKO clones and Clone F2 was selected for further experiments.

### FLIPR assay to measure constitutive Ca^2+^ entry

HEK293T (ATCC^®^ CRL-3216^™^) and S1/S2 DKO HEK293 cells were maintained in cell culture in 1X DMEM (41965-039, Thermo Fisher Scientific) supplemented with 10% FBS, 1 mM sodium pyruvate, 10 mM HEPES, 1X MEM non-essential amino acids and 1% Penicillin-Streptomycin. The FLIPR assay to measure the constitutive Ca^2+^ entry was adapted from our previous study (Bulla et al., 2019). Briefly, the cells were seeded in Corning® 96-well black polystyrene clear bottom microplates (CLS3603, Sigma–Aldrich) at a density of 30,000 in 100 µl medium per well. The medium contained ∼0.2 mM CaCl_2_ to prevent excessive constitutive Ca^2+^ entry after transfection of constitutively active constructs. Since FBS is estimated to contain 3.5 – 4 mM Ca^2+^ (Blankenship & Heitman, 2005), low Ca^2+^ (∼0.2 mM) containing medium was formulated by supplementing the Ca^2+^ free DMEM (21068028, Thermo Fisher Scientific) with 6% FBS. Other supplements were 1 mM sodium pyruvate, 10 mM HEPES and 1X MEM non-essential amino acids. A total of 200 ng plasmid DNA was transfected per well using Lipofectamine^®^ 2000 (Thermo Fisher Scientific). The growth medium of the cells was removed 16–20 h after transfection and the cells were loaded with 50 μl of Calcium 5 indicator (FLIPR^®^ Calcium 5 assay kit, R8186, Molecular Devices) prepared in modified Krebs buffer containing 0.2 mM CaCl_2_, 1 mM MgCl_2_,140 mM NaCl, 4.8 mM KCl, 10 mM D-glucose and 10 mM HEPES (pH 7.4). Cells were incubated in the loading buffer at 37°C for 1 h and were excited using a 470–495 nm LED module of the FLIPR, and the emitted fluorescence signal was filtered with a 515–575 nm emission filter. After recording a 50 s baseline, 50 μl of 3.8 mM CaCl_2_-containing Krebs buffer was administered to the cells, resulting in 2 mM final concentration of Ca^2+^. The changes in fluorescence intensity were measured for first 60 s after CaCl_2_ administration with an acquisition rate of 2 Hz and for further 180 s with 0.45 Hz. The fluorescence signals were analyzed using the FLIPR Tetra software, ScreenWorks 3.1.1.8 (Molecular Devices). To quantify the constitutive Ca^2+^ entry dedicated to each mutant, a ratio of area under the curve (AUC) of the Ca^2+^ entry traces of the mutant Orai to that of wild type was calculated.

### Electrophysiology

Whole-cell patch-clamp experiments were specifically designed to measure two distinct types of inward Ca^2+^ currents:

i. Canonical *I*_CRAC_ evoked by IP_3_ and facilitated by interacting hO1 and hSTIM1 molecules.
ii. Constitutive Ca^2+^ influx mediated by the selected constitutively active hO1 mutants in the absence of hSTIM1 and IP_3_.

#### I_CRAC_ assay

The experimental procedure was described in detail previously (Bhardwaj et al., 2020). Briefly, HEK293 S1/S2 DKO cells were trypsinized and seeded into the 6-well plate. When cells reached confluency of approx. 70-80%, they were transiently co-transfected with plasmids encoding WT hO1 and WT STIM1. 5 μl Lipofectamine^®^ 2000 (Thermo Fisher Scientific) and 2+2 μg of plasmid DNA encoding the proteins were mixed in Opti-MEM^™^ (Thermo Fisher Scientific) and applied onto the cells in a single well. After 6 h of incubation with the Lipofectamine-DNA complexes, cells were trypsinized and reseeded sparsely into the patch-clamp-compatible 35 x 10 mm cell culture petri dishes.

*I*_CRAC_ was measured after further 18 h of incubation at 37 °C in humidified 5% CO_2_ atmosphere (24 h in total after transfection). Only cells exhibiting comparable (modest) fluorescence levels of both mCherry-Orai1 and GFP-STIM1 were selected for recordings. The experiments were performed at room temperature, in whole-cell configuration. Pipettes were pulled from 1.5 mm thin-wall borosilicate glass capillaries with filament (BF150-86-7.5, Sutter Instruments) using a horizontal P-1000 puller (Sutter Instruments) to obtain a serial resistance of around 2.5 MΩ. Currents were recorded with PatchMaster software (HEKA Elektronik), using an EPC-10 USB amplifier (HEKA Elektronik). Upon establishment of giga seal and successful break-in into the single, mCherry fluorescent cell, 50 ms voltage ramps spanning −150 to +150 mV were delivered from a holding potential of 0 mV every 2 seconds. Currents were filtered at 2.9 kHz and digitized. Liquid junction potential was 10 mV and currents were determined and corrected before each voltage ramp. Leak currents were corrected by subtracting the initial ramp currents from all subsequent currents using FitMaster software (HEKA Elektronik). Currents were extracted at −80 and +80 mV and normalized to cell capacity (size).

At break in, the bath solution contained 120 mM NaCl, 10 mM tetraethylammonium chloride (TEA-Cl), 2 mM MgCl_2_, 10 mM CaCl_2_, 10 mM HEPES, and 32 mM glucose (pH 7.2 with NaOH, 300 mOsmol). After 60 seconds of recording, cells were perfused with nominal Ca^2+^-free bath solution (the same as above, but without 10 mM CaCl_2_, osmolarity was adjusted with more glucose).

Calcium-free internal solution contained 120 mM Cs-glutamate, 3 mM MgCl_2_, 10 mM HEPES, 0.05 mM D-myo-inositol 1,4,5-trisphosphate, trisodium salt (IP_3_) (407137, Calbiochem) and 20 mM EGTA (pH 7.2 with CsOH, 310 mOsmol with glucose).

#### Constitutive Ca^2+^ influx assay

This experimental approach was largely identical to the *I*_CRAC_ assay, with two significant modifications:

i. HEK293 S1/S2 DKO cells were transfected each time only with a single plasmid encoding the relevant hO1 mutant. Unlike in the previous assay, no STIM1 was co-expressed.
ii. Calcium-free internal solution contained 120 mM Cs-glutamate, 3 mM MgCl_2_, 10 mM HEPES and 20 mM BAPTA (B1212, Invitrogen), not EGTA (pH 7.2 with CsOH, 310 mOsmol adjusted with glucose). No IP_3_ was added to the solution.

### NFAT1 translocation assay

Fluorescence microscopy and confocal NFAT subcellular localization studies were performed as described earlier (Schober et al., 2019). ImageJ was used to analyze subcellular NFAT localization by intensity measurements of the cytosol and the nucleus (nucleus/cytosol ratios: inactive (<0.85), homogenous (0.85–1.15), and active (>1.15)). All represented images of Orai isoforms as well as NFAT localization were created with a custom-made software integrated into MATLAB (v7.11.0, The MathWorks, Inc.). The experiments were performed on three different days at room temperature and the resulting data of all experiments are presented as mean ± S.D. (standard deviation) for the indicated number of experiments.

### Cell Surface Biotinylation

Cell surface biotinylation was performed as described (Simonin & Fuster, 2010). Cells were rinsed with 1x PBS and surface proteins were biotinylated by incubating cells with 1.5 mg/ml sulfo-NHS-LC-biotin in 10 mM triethanolamine (pH 7.4), 1 mM MgCl_2_, 2 mM CaCl_2_, and 150 mM NaCl for 90 min with horizontal motion at 4°C. After labeling, plates were washed with quenching buffer (PBS containing 1 mM MgCl_2_, 0.1 mM CaCl_2_, and 100 mM glycine) for 20 min at 4°C, then rinsed once with 1X PBS. Cells were then lysed in RIPA buffer [150 mM NaCl, 50 mM Tris·HCl (pH 7.4), 5 mM EDTA, 1% Triton X-100, 0.5% deoxycholate, and 0.1% SDS] and lysates were cleared by centrifugation. Cell lysates of equivalent amounts of protein were equilibrated overnight with streptavidin agarose beads at 4°C. Beads were washed sequentially with solutions A [50 mM Tris·HCl (pH 7.4), 100 mM NaCl, and 5 mM EDTA] three times, B [50 mM Tris·HCl (pH 7.4) and 500 mM NaCl] two times, and C (50 mM Tris·HCl, pH 7.4) once. Biotinylated proteins were then released by heating to 95°C with 2.5X Lämmli buffer.

### Structural modeling

The structural model of hO1 was generated using Modeller 9.21 (Fiser, Do, & Sali, 2000; Sali & Blundell, 1993; Webb & Sali, 2016) based on the closed-latched structure (PDB ID: 4HKR (Hou et al., 2012)), downloaded from the Orientations of Proteins in Membranes (OPM) database (Lomize, Lomize, Pogozheva, & Mosberg, 2006). Sequences of the template and query proteins were aligned using ClustalW 2.1 (Larkin et al., 2007). During model optimization, main chain atoms of residues 72-106, 121-143, and 235-291, roughly corresponding to the TM regions, were kept fixed in order to avoid large conformational changes. Six-fold symmetry constraints were introduced for main chain atoms of residues 99-128, 138-178, and 190-242, corresponding to loop regions and flanking TM regions. Based on PSIPRED 4.0 (Buchan, Minneci, Nugent, Bryson, & Jones, 2013; Jones, 1999), α-helical constraints were introduced on residues 117-154, 164-199, and 231-286. Loop modeling protocol was applied on residues 107-120, 144-164, and 198-234. In total 50 models were built, and 10 loop modeling tries were attempted on each model. The final model was chosen based on lowest objective function values as reported by Modeller for loop modeling, and visual inspection to avoid protein loops protruding into the anticipated position of the membrane bilayer.

### Molecular dynamics (MD) simulations

Simulation systems were prepared and mutations were introduced using CHARMM-GUI (Brooks et al., 2009; Jo et al., 2014; Jo, Kim, & Im, 2007; Jo, Kim, Iyer, & Im, 2008; Jo, Lim, Klauda, & Im, 2009; Lee et al., 2016; Wu et al., 2014), E190 was protonated, and truncated termini were acetylated/methylated. Two Ca^2+^ ions were introduced in the structure, one at the position determined by X-ray crystallography in the structure (PDB ID: 4HKR), and another 5 Å below along the axis of the pore. The protein was surrounded by palmitoyl-oleoyl-phosphatidylcholine (POPC) lipids in a rectangular box with size of 120.12×120.12×129.76 Å and solvated in TIP3 water with neutralizing ions and 150 mM NaCl. The simulations were performed using GPU-accelerated *pmemd* of AMBER 18 (Case et al., 2018) using the CHARMM36m force-field (Huang et al., 2017), under NPγT conditions, 1 bar pressure, constant surface tension of zero and a temperature of 310 K. Equilibration of simulation systems was performed as per the protocol prescribed by CHARMM-GUI. Motions of protein variants were simulated for 200 ns for 5 independent replicas each.

### Analysis of MD trajectories

Contact analysis was performed using the *analysis.distances.contact_matrix* function of MDAnalysis (Michaud-Agrawal, Denning, Woolf, & Beckstein, 2011) (Gowers et al., 2016) on all atoms of residues, with a cut-off range of 5 Å. Inter- and intra-subunit atomic contacts in the Orai hexamer have been distinguished and counted separately. Residues were taken to be in contact if any pair of their atoms were in contact. Helicity analysis was performed by custom scripts based on MDAnalysis to calculate for each residue *i* the angle *α_i_* = *ψ_*i*_* + *φ_i_*_+1_, where *φ_*i*_* and *ψ_i_* are the Ramachandran-angles for residue *i*. For residues in a perfect α-helix, the value of *α* = −105°. During helicity analysis, odd (A, C, E chains with a more extended TM4-TM4ext region) and even (B, D, F chains with a helix-turn-helix/backfolded TM4-TM4ext region) subunits of the Orai hexamer were distinguished owing to the 3-fold symmetry of the starting structure. Pore diameter analysis was performed by HOLE using elements of the programming interface from MDAnalysis (Smart, Goodfellow, & Wallace, 1993; Smart, Neduvelil, Wang, Wallace, & Sansom, 1996). Pore diameters were calculated every ns of the simulations and averaged over all frames.

## Acknowledgements

Calculations were performed on UBELIX (http://www.id.unibe.ch/hpc), the HPC cluster at the University of Bern. We would like to thank Dr. Stefan Mueller, Thomas Schaffer and Bernadette Nyfeler at the FACS facility, Institute of Pathology, University of Bern for their help in FACS sorting of CRISPR/Cas9-generated knockout cells. Also, we thank Tamara Locher, University of Bern for her technical assistance in cell culture. We thank Prof. Francisco Javier Martin-Romero, University of Extremadura, Spain for providing us the mO1-eGFP construct, Dr. Matthias Seedorf, Heidelberg University, Germany for providing us the mCherry-tagged hO1 and hO3 constructs and Prof. Christoph Romanin, University of Linz, Austria for providing us the CFP-hO2 construct. We are thankful to Dr. Anant B. Parekh, National Institute of Environmental Health Sciences, North Carolina, USA for providing important feedback on our manuscript. H.G. thanks Austrian Science Fund (FWF) for PhD scholarship through W1250 NanoCell PhD program. I.F. was funded by FWF project P32075.-B. B.A., G.G., M.A.H. and R.B. were funded by the Swiss National Science Foundation Sinergia grants (CRS115_180326 and CRS113_160782). R.B. was also supported by the Marie Curie Actions International Fellowship Program IFP TransCure, University of Bern, Switzerland (from 2014 to 2017).

## Additional information

### Funding

| Funder | Grant reference number | Author |
| --- | --- | --- |
| Swiss National Science Foundation | CRS115_180326 | Matthias A. Hediger |
| Swiss National Science Foundation | CRS113_160782 | Matthias A. Hediger |
| Austrian Science Fund | P32075.-B | Irene Frischauf |
| The funders had no role in study design, data collection and interpretation, or the decision to submit the work for publication. |  |  |

## Author contributions

Bartlomiej Augustynek, Electrophysiology Experimentation, Data Analysis and Curation, Writing - original draft, Writing - review and editing; Gergely Gyimesi, Homology modeling, Molecular dynamics simulations, Data Analysis, Writing - original draft, Writing - review and editing; Jan Dernič, FLIPR Experimentation; Matthias Sallinger, NFAT1 translocation Experimentation; Giuseppe Albano, Cell surface biotinylation Experimentation; Gabriel Jonathan Klesse, Site-directed mutagenesis; Palanivel Kandasamy, CRISPR/Cas9 knockout generation; Herwig Grabmayr, NFAT1 translocation Experimentation; Irene Frischauf, Writing - original draft, Writing - review and editing; Daniel G. Fuster, Supervision, Methodology, Writing – review and editing; Christine Peinelt, Supervision, Methodology, Writing - review and editing; Matthias A. Hediger, Project Strategy, Project Coordination and Supervision, Manuscript Writing; Rajesh Bhardwaj, Conceptualization, Supervision, Validation, Investigation, Site-directed mutagenesis and FLIPR Experimentation, CRISPR/Cas9 knockout generation, Data Analysis and Curation, Visualization, Methodology, Writing - original draft, Project administration, Writing - review and editing.

## Supplementary Figures

**Figure 2- figure supplement 1.**
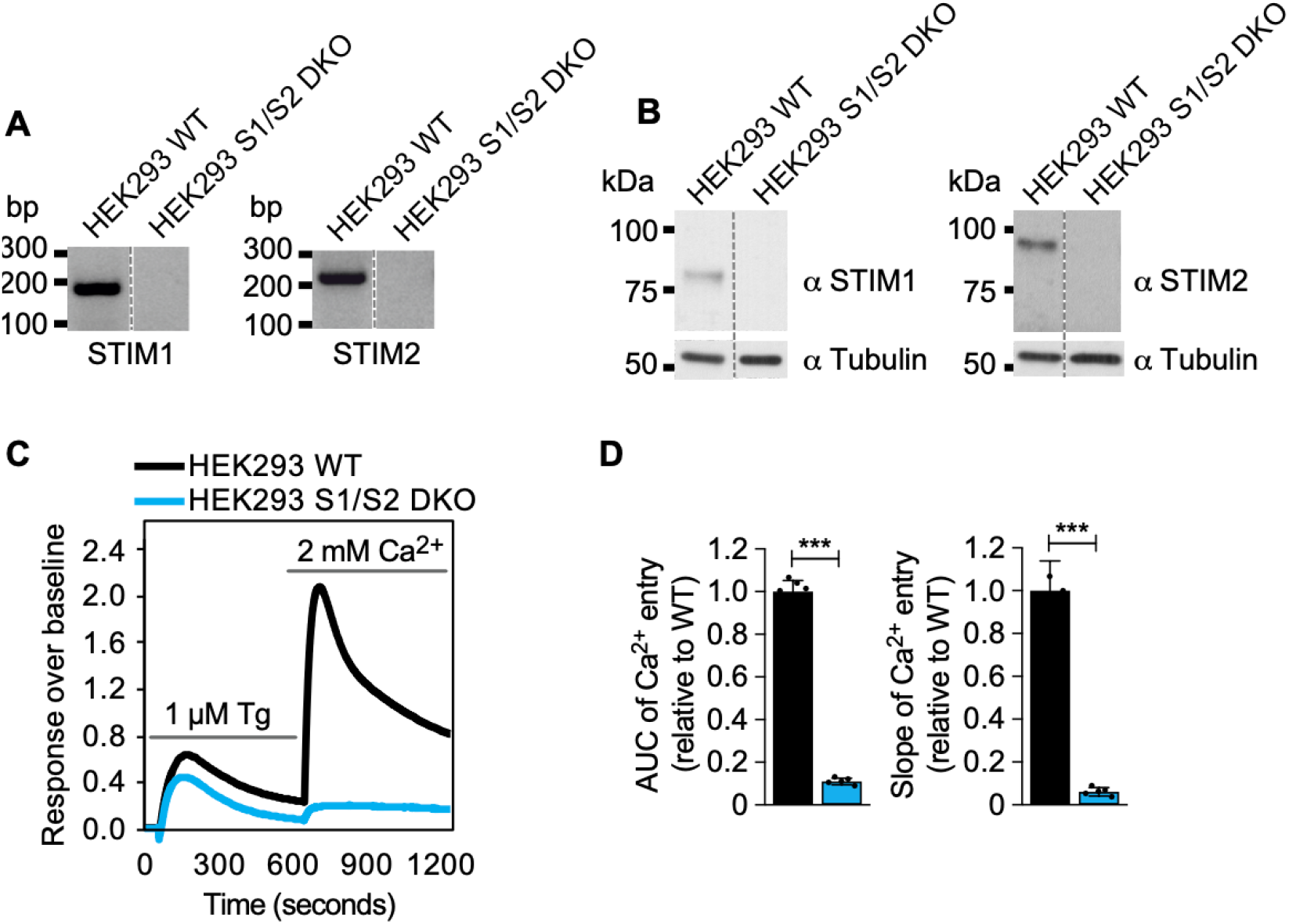
Validation of HEK293 S1/S2 DKO cells. (**A**) Agarose gel electrophoresis showing PCR products of STIM1 and STIM2 using genomic DNA template isolated from HEK293 WT and S1/S2 DKO cells. (**B**) Western blot analysis of HEK293 WT and S1/S2 DKO cells using human STIM1 and STIM2 antibodies. (**C**) Representative SOCE measurement traces from HEK293 WT and S1/S2 DKO cells treated with 1 µM thapsigargin (Tg) in nominally calcium free buffer followed by add-back of 2mM CaCl_2_. (**D**) Quantifications of area under the curve (AUC) and slope of the Ca^2+^ entry traces (second peak in “C”) represented as mean ±SD; n=5.

**Figure 2- figure supplement 2.**
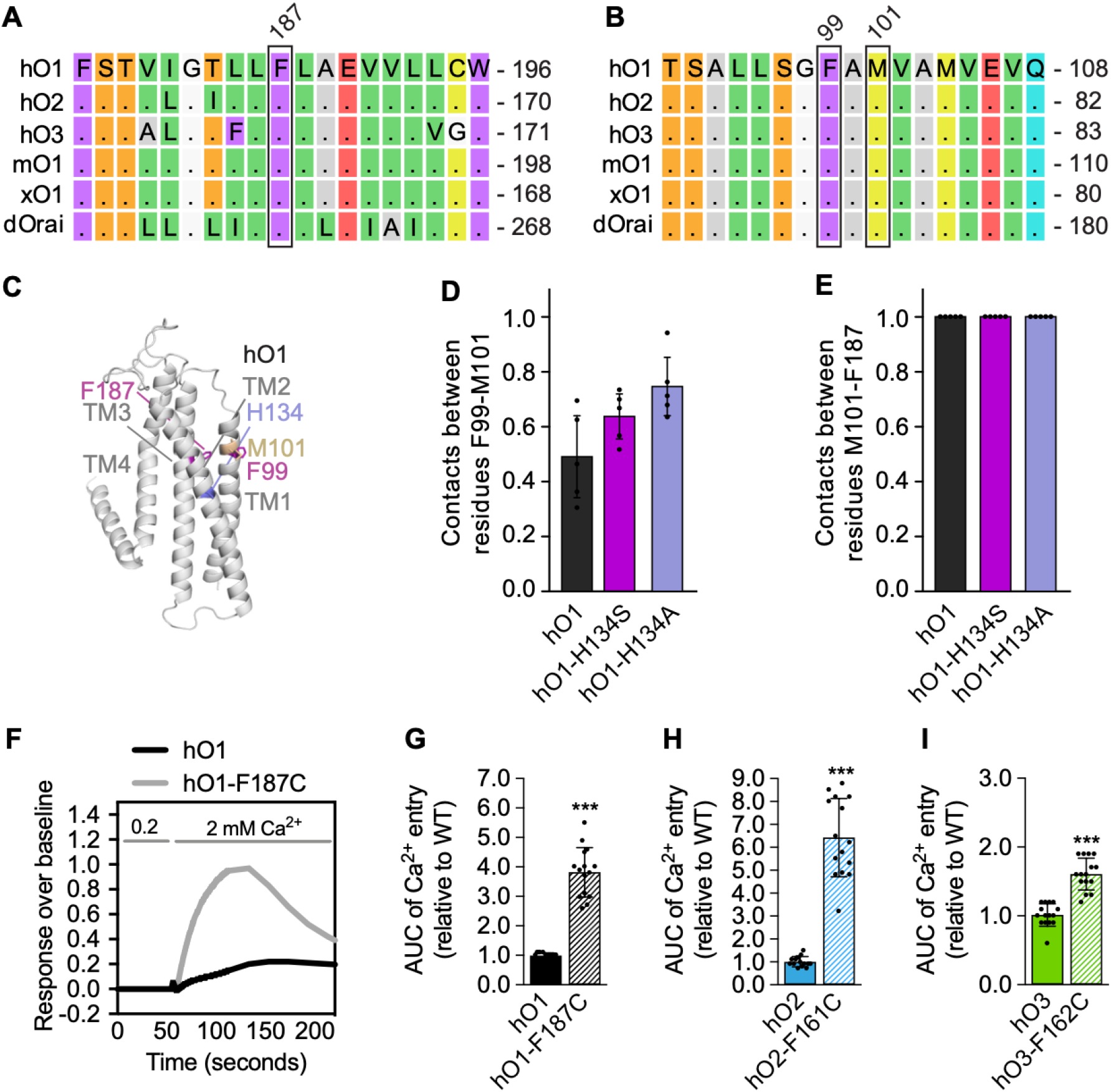
Inter-subunit F99-M101 contacts in hO1 WT and H134 mutant, and constitutive activity of hO1 TM3 F187C mutant and corresponding hO2 and hO3 mutants. (**A**) Multiple sequence alignment of TM3 of hO1, hO2, hO3, mO1, xO1 and dOrai is shown, highlighting the conserved F187 residue. (**B**) Multiple sequence alignment of TM1 of hO1, hO2, hO3, mO1, xO1 and dOrai is shown, highlighting the conserved F99 and M101 residue. (**C**) Cartoon representation of a single subunit of hO1 model depicting indicated residues. (**D**) Frequencies of contacts between F99 and M101 in hO1 WT, H134S and H134A mutant channels (mean ± SD; n=5). (**E**) Contact frequencies between M101 and F187 in WT hO1, hO1-H134S and hO1-H134A mutant channels (mean ± SD; n=5). (**F**) Representative constitutive Ca^2+^ entry traces of HEK293 cells transfected with WT hO1 and hO1-F187C constructs with initial baseline recording in 0.2 mM CaCl_2_, followed by addition of 2 mM CaCl_2_. The quantified AUC of Ca^2+^ entry peak from HEK293 cells expressing (**G**) WT hO1 and hO1-F187C, (**H**) WT hO2 and hO2-F161C and (**I**) WT hO3 and hO3-H162C (mean ± SD; n=15). *p* ≤ 0.001 is indicated as “***”.

**Figure 3- figure supplement 1.**
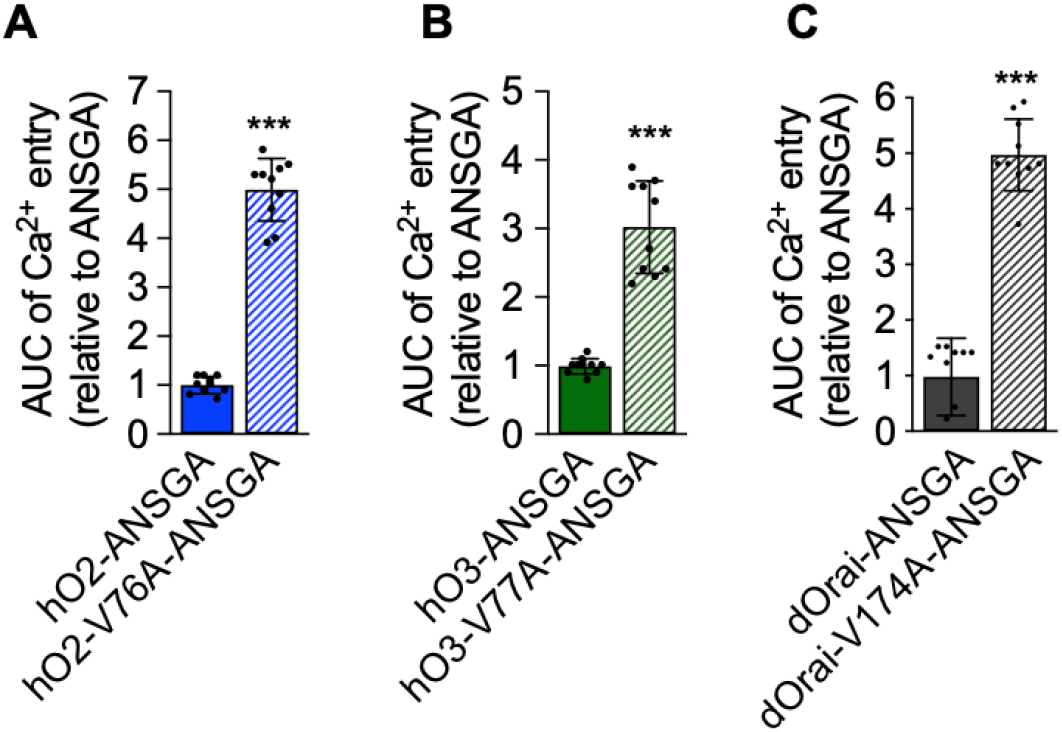
Insertion of pore hydrophobic gate (Val) mutations in the ANSGA mutants of hO2, hO3 and dOrai leads to their constitutive activation. (**A-C**) AUC of the constitutive Ca^2+^ entry traces from indicated Orai mutants expressed in HEK293 S1/S2 DKO cells (mean ± SD; n=10). *P* ≤ 0.001 is indicated as “***”.

**Figure 4- figure supplement 1.**
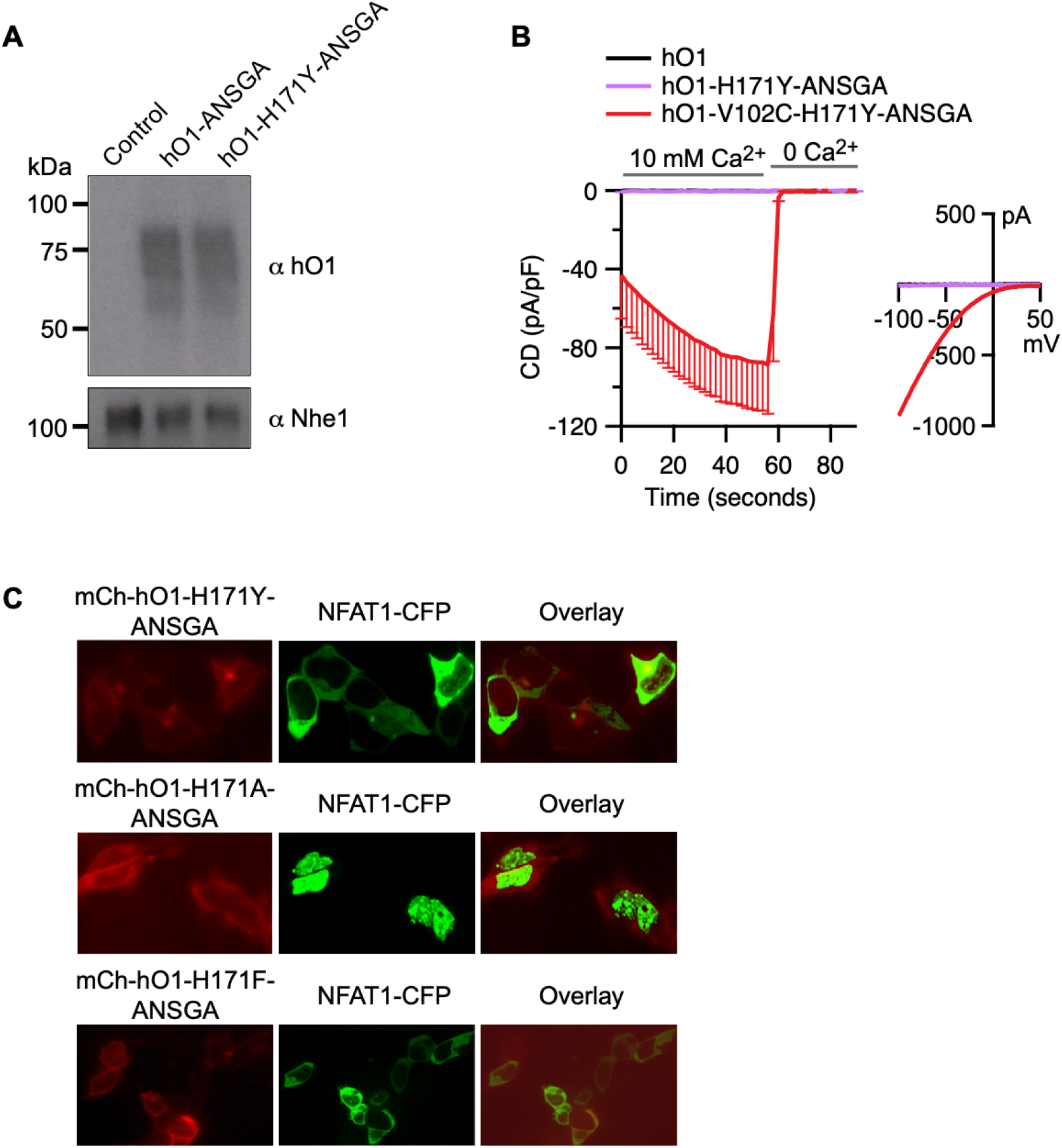
H171Y substitution inhibits the constitutive activity of hO1-ANGSA channel without affecting the plasma membrane localization and assembly of the channel. (**A**) hO1 immunoblot of surface biotinylated mCherry control, mCherry-hO1-ANSGA and mCherry-hO1-H171Y-ANSGA proteins expressed in HEK293 S1/S2 DKO cells. Plasma membrane protein Nhe1 is shown as a loading control. (**B**) Current densities (CD) of the constitutive Ca^2+^ currents recorded from HEK293 S1/S2 DKO cells transiently overexpressing: WT hO1 (n=6), hO1-H171Y-ANSGA (n=8) and hO1-V102C-H171Y-ANSGA (n=8), presented as average values, -SEM, with corresponding average current-voltage (I/V) relations extracted at t= 59s. (**C**) Representative confocal microscopy images of HEK293 cells co-expressing NFAT1-CFP with either mCherry-hO1-H171Y-ANSGA, hO1-H171A-ANSGA or hO1-H171F-ANSGA constructs along with the CFP/mCherry overlay images.

**Figure 5- figure supplement 1.**
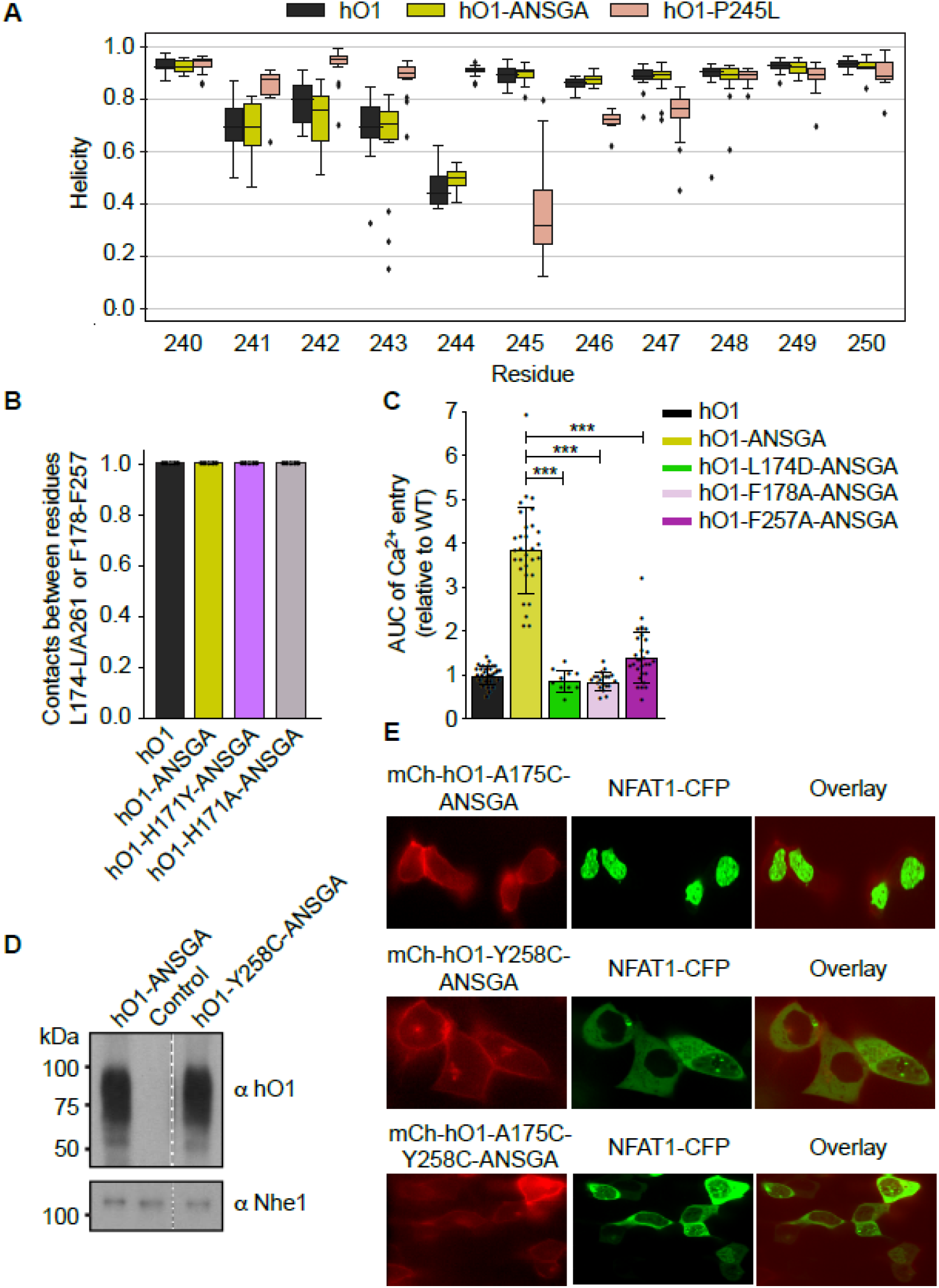
Relevance of selected TM3 and TM4 residues in the constitutive activity of hO1-ANSGA channel. (**A**) Helicity of TM4 residues around position 245 of hO1 averaged over 5 MD trajectories for chains A, C, and E, giving 15 points per residue and system (see Methods). Box-and-whiskers plot showing quartiles is used, outliers are plotted individually. (**B**) Intra-subunit contact frequencies between residue pairs L174-L/A261 and F178-F257 averaged over each of the 5 MD trajectories are shown for various simulation systems. Data are identical for both residue pairs. (**C**) The AUC of constitutive Ca^2+^ entry recorded in HEK293 S1/S2 DKO cells expressing mCherry-tagged WT hO1 or indicated ANSGA variants (mean ± SD; n ≥ 10). *p* ≤ 0.001 is indicated as “***”. (**D**) hO1 immunoblot of surface biotinylated mCherry control, mCherry-hO1-ANSGA and mCherry-hO1-Y258C-ANSGA proteins expressed in HEK293 S1/S2 DKO cells along with plasma membrane protein Nhe1 shown as a loading control. (**E**) Representative confocal microscopy images of HEK293 cells co-expressing NFAT1-CFP and either of the indicated mCherry-Orai-ANSGA constructs along with the CFP/mCherry overlay images.

**Figure 5- figure supplement 2.**
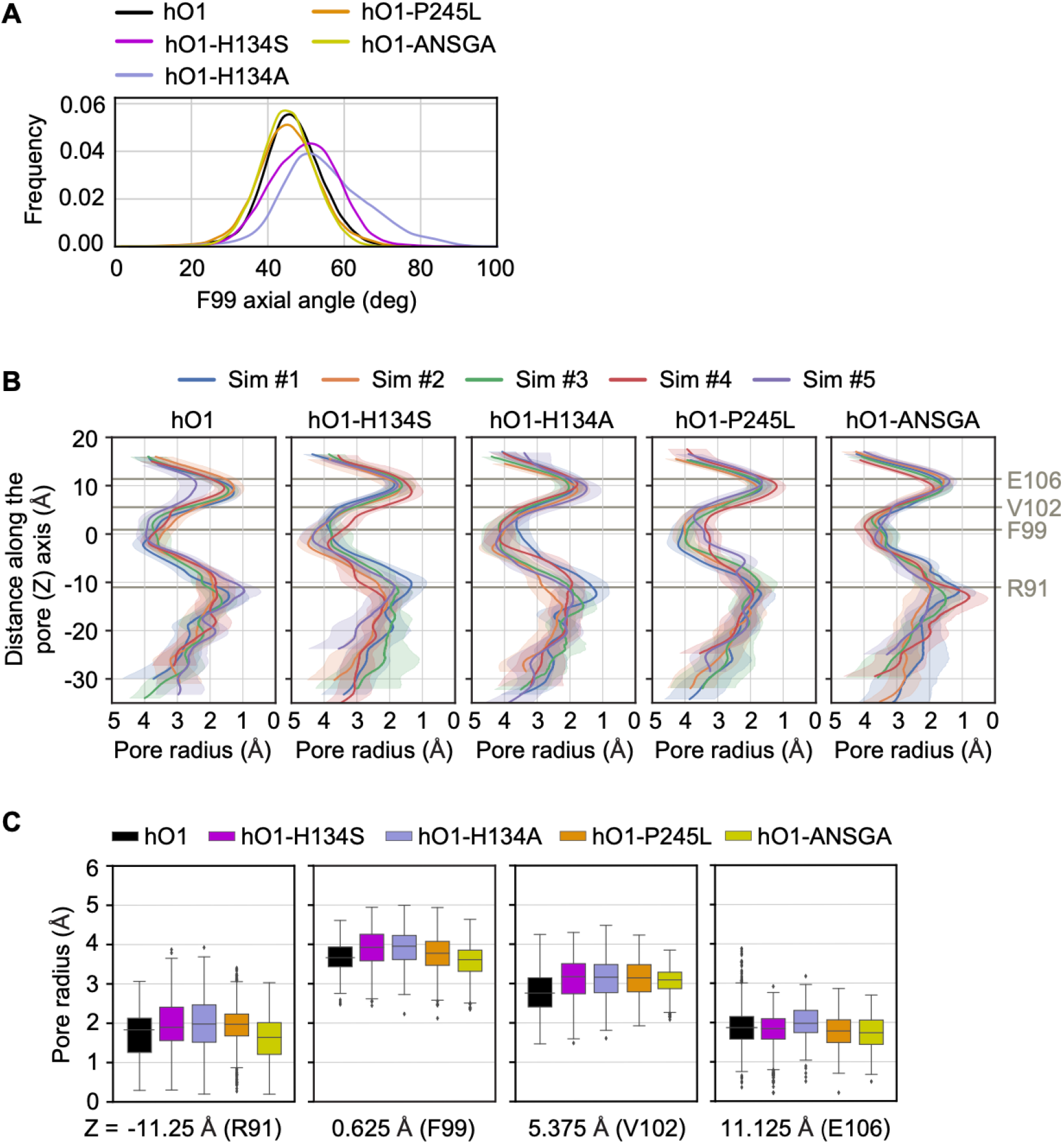
Impact of H134S, H134A, P245L and ANSGA mutations on TM1 rotation and pore dilation. (**A**) Axial angle distribution of the F99 side-chain of hO1 as defined by Yamashita *et al*., 2017 are shown (Yamashita et al., 2017) for WT hO1, hO1-H134S, hO1-H134A, hO1-P245L and hO1-ANSGA mutants. In loose terms, the axial angle measures the angle between the pore axis, the center of mass of the C_α_ atoms of residues 96-102, and the C_α_ atom of residue 99. Values are averaged for all 5 MD trajectories and for all six subunits of the Orai hexamer. (**B**) Pore radius as calculated by the HOLE program for each MD trajectory of various simulation systems. Shaded regions show average and SD of pore radius. Residues of functional importance in TM1 are marked for scale. (**C**) For comparison, values of pore radius along the trajectory of the Orai channel variants at cross-sections corresponding to the positions of TM1 residues marked in panel B are shown. For each system and cross-section, pore radii for 200 frames from 5 simulations (total 1000 points) are plotted using a box-and-whiskers plot showing quartiles and outliers.

**Figure 6- figure supplement 1.**
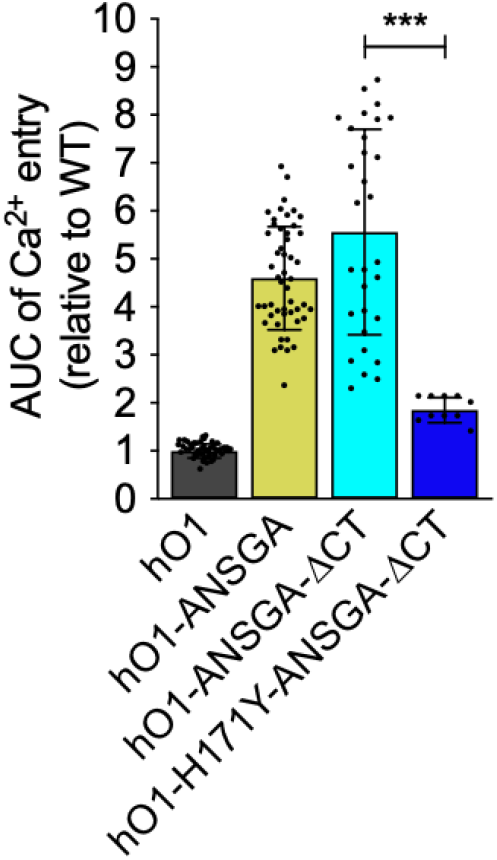
The TM4 extension beyond the ANSGA region of hO1 is dispensable for H171Y mediated inhibition of constitutive activity. The quantified AUC of constitutive Ca^2+^ entry recorded in HEK293 S1/S2 DKO cells expressing the indicated hO1-ANSGA mutants relative to the WT hO1 channel (mean ± SD; n=10). *p* ≤ 0.001 is indicated as “***”.

**Figure 7- figure supplement 1.**
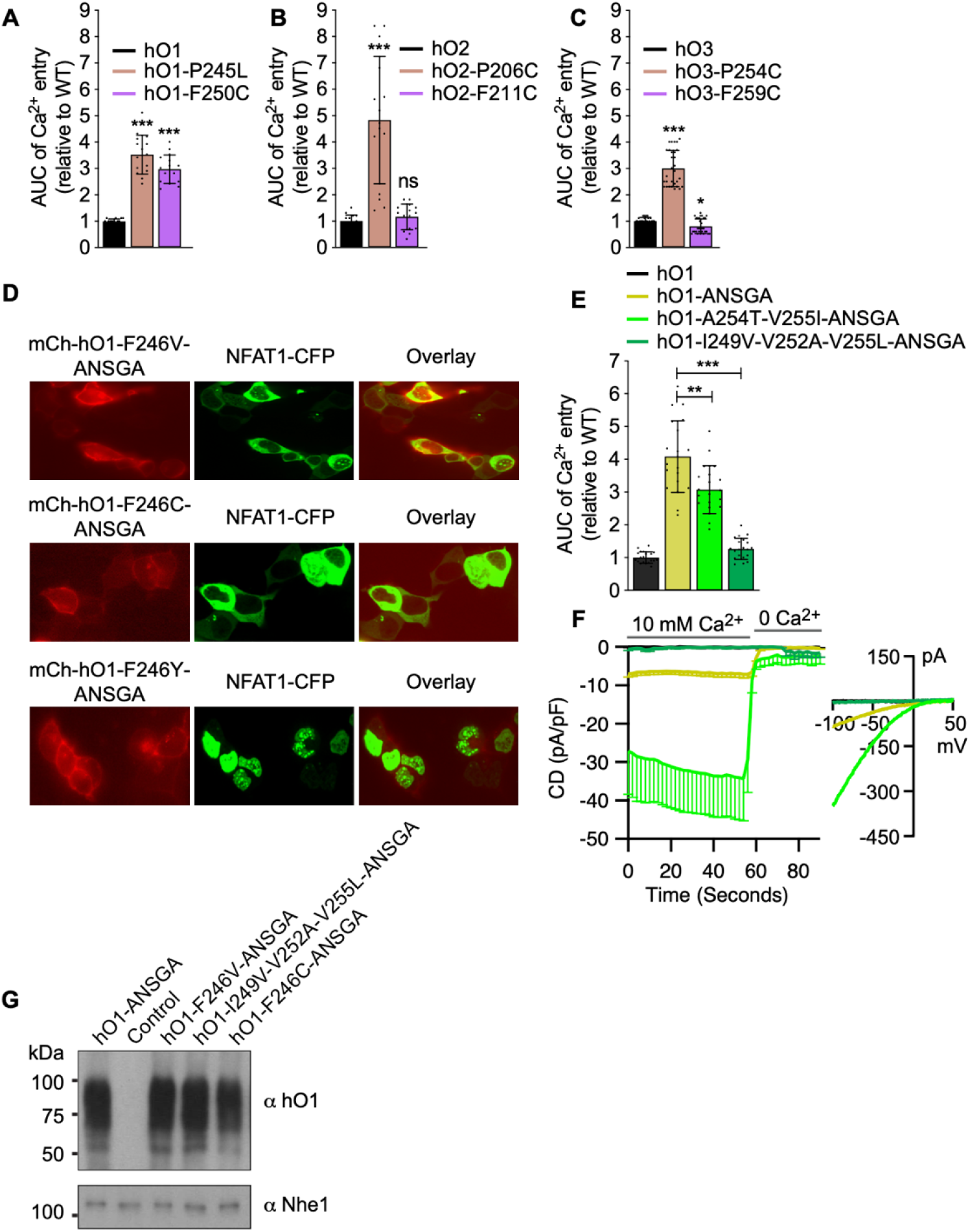
The quantified AUC of constitutive Ca^2+^ entry recorded in HEK293 cells expressing (**A**) hO1-P245L and hO1-F250C, (**B**) hO2-P206C and hO2-F211C and (**C**) hO3-P254C and hO3-F259C relative to their respective WT Orais (mean ± SD; n ≥ 15). *p* ≤ 0.001 is indicated as “***”, 0.01 < *p* < 0.05 as “*” and *p* ≥ 0.05 as “ns”. (**D**) Representative confocal microscopy images of HEK293 cells co-expressing NFAT1-CFP with either F246V, F246C or F246Y mutants of mCherry-hO1-ANSGA along with the CFP/mCherry overlay images. (**E**) AUC quantifications of constitutive Ca^2+^ influx measured in HEK293 S1/S2 DKO cells expressing hO1-ANSGA or its A254T-V255I and I249V-V252A-V255L mutants relative to WT hO1 (mean ± SD; n=20). *p* ≤ 0.001 is indicated as “***” and 0.001 < *p* ≤ 0.01 as “**”. (**F**) Current densities (CD) of the constitutive Ca^2+^ currents recorded from HEK293 S1/S2 DKO cells transiently overexpressing: WT hO1 (n=6), hO1-ANSGA (n=23), hO1-A254T-V255I-ANSGA (n=8) and hO1-I249V-V252A-V255L-ANSGA (n=6), presented as average values, -SEM, with corresponding average current-voltage (I/V) relations extracted at t= 59s. (**G**) hO1 immunoblot of surface biotinylated mCherry control, mCherry-hO1-ANSGA and other indicated ANSGA mutant proteins expressed in HEK293 S1/S2 DKO cells. Plasma membrane protein Nhe1 is shown as a loading control.

**Figure 7- figure supplement 2.**
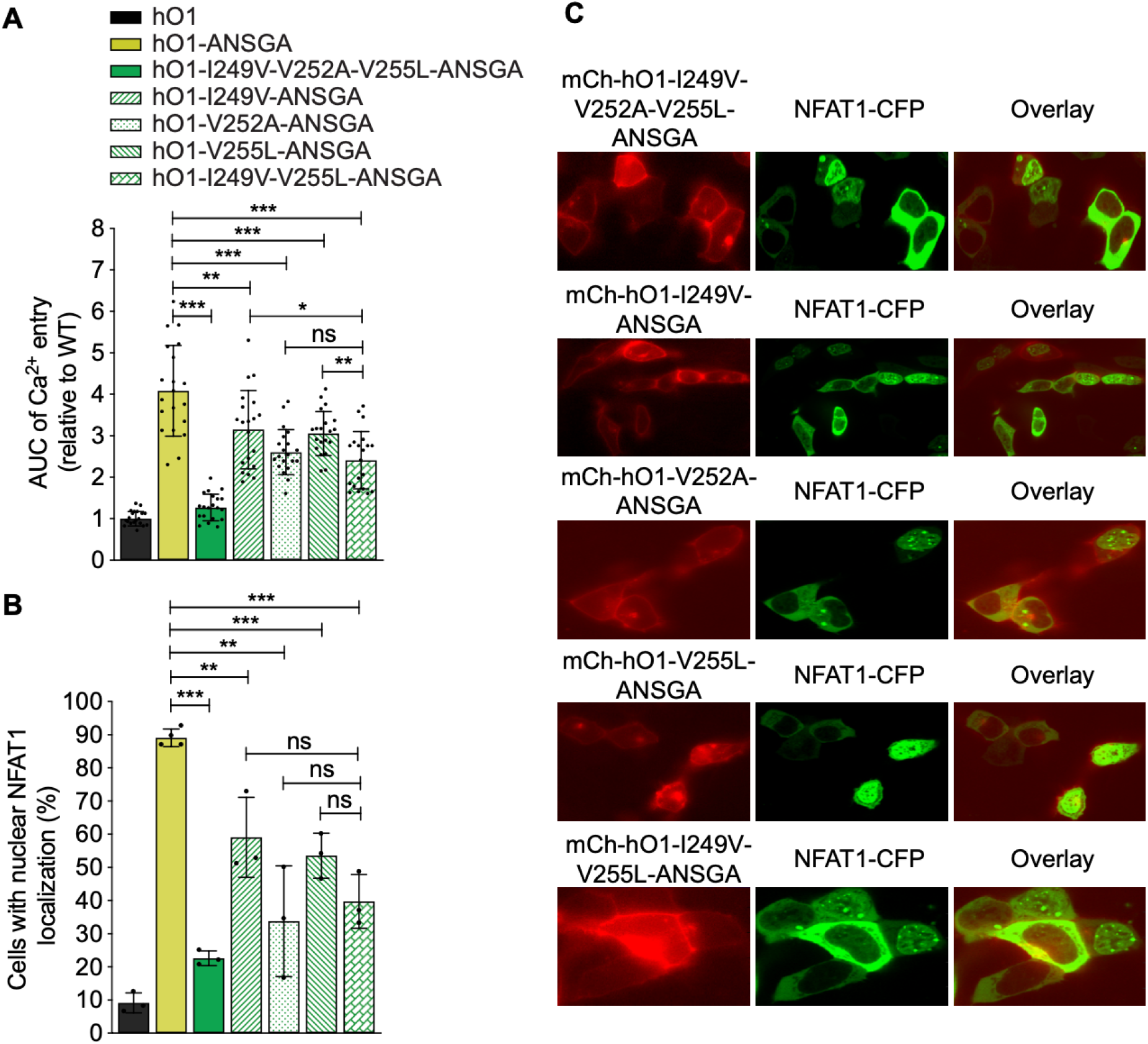
(**A**) AUC quantifications of constitutive Ca^2+^ influx measured in HEK293 S1/S2 DKO cells expressing hO1-ANSGA or other indicated mutants of hO1-ANSGA relative to WT hO1 (mean ± SD; n=20). (**B**) HEK293 cells expressing mCherry-hO1, hO1-ANSGA or I249V-V252A-V255L, I249V, V252A, V255L and I249V-V255L mutants of hO1-ANSGA with nuclear NFAT1-CFP localization shown as percentage (mean ± SD; n ≥ 3). *p* ≤ 0.001 is indicated as “***”, 0.001 < *p* ≤ 0.01 as “**”, 0.01 < *p* < 0.05 as “*” and *p* ≥ 0.05 as “ns”. (**C**) Representative confocal microscopy images of HEK293 cells co-expressing NFAT1-CFP with either mCherry-hO1, mCherry-hO1-ANSGA or indicated mutants of mCherry-hO1-ANSGA along with the CFP/mCherry overlay images.

**Figure 8- figure supplement 1.**
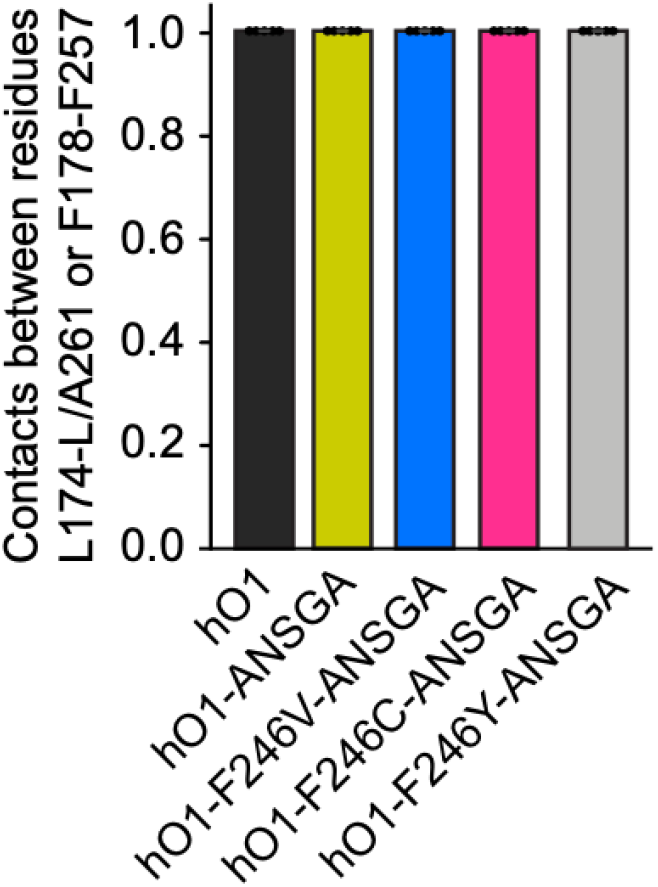
Intra-subunit contact frequencies between residue pairs L174-L/A261 and F178-F257 averaged over each of the 5 MD trajectories are shown for various simulation systems. Data are identical for both residue pairs.

**Figure 9- figure supplement 1.**
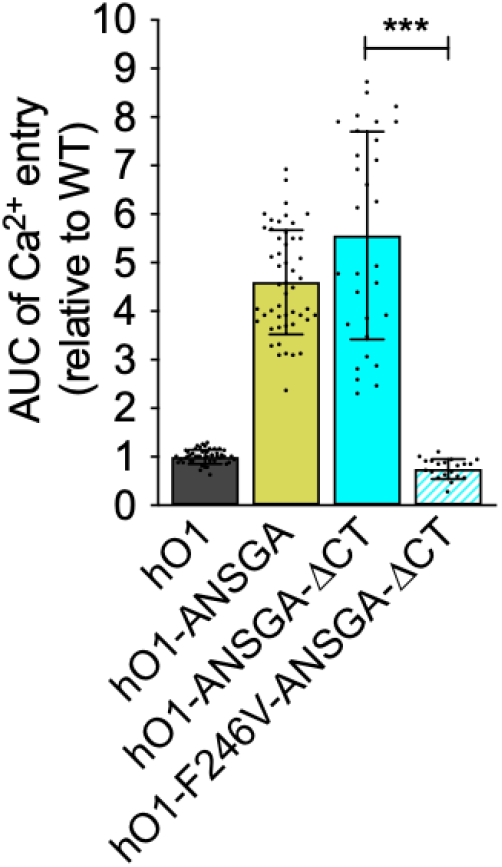
The TM4 extension beyond the ANSGA region of hO1 is dispensable for F246V mediated inhibition of constitutive activity. The quantified AUC of constitutive Ca^2+^ entry recorded in HEK293 S1/S2 DKO cells expressing the indicated hO1-ANSGA mutants relative to the WT hO1 channel (mean ± SD; n=10). *p* ≤ 0.001 is indicated as “***”.

## Key resources table

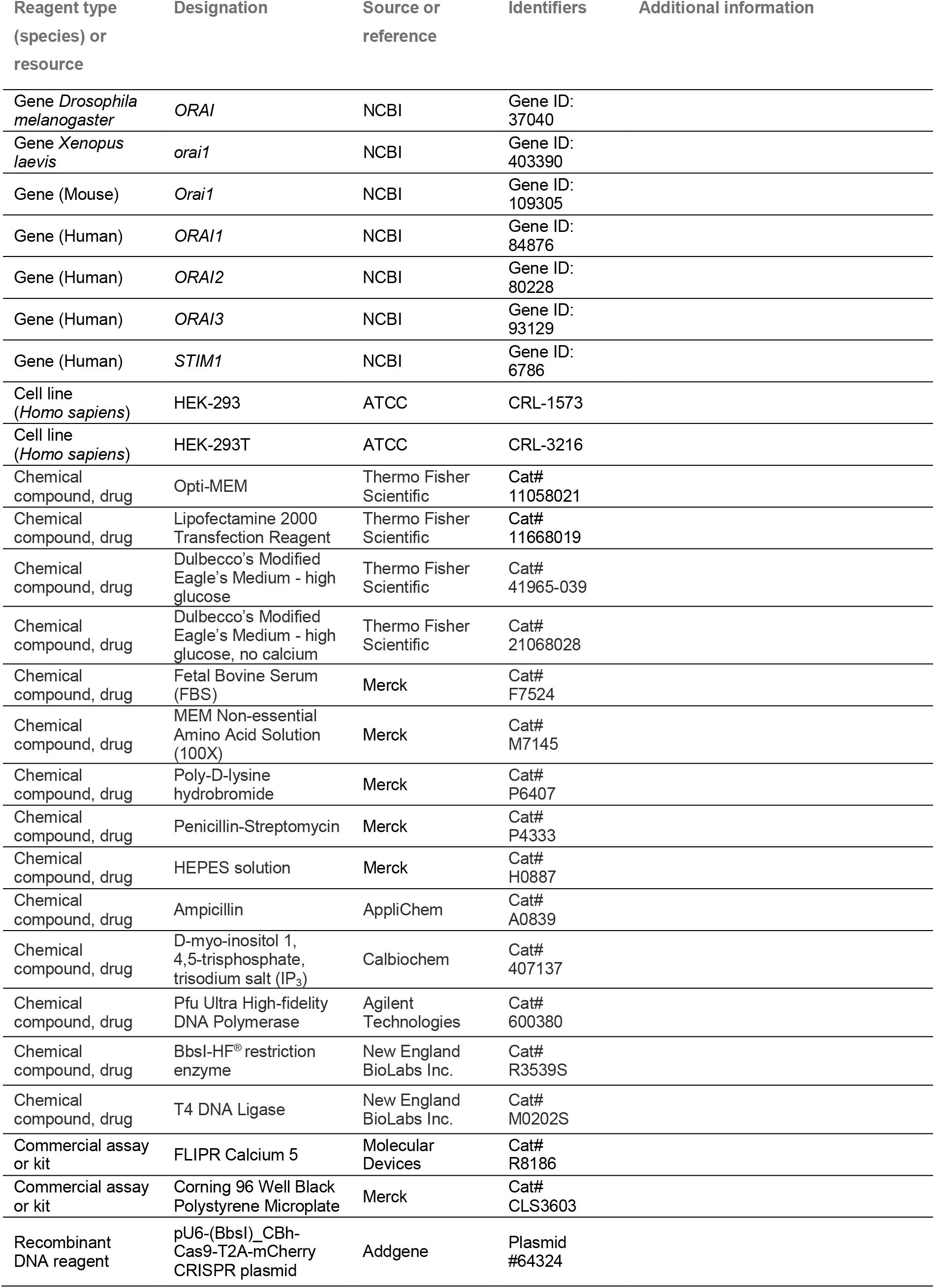
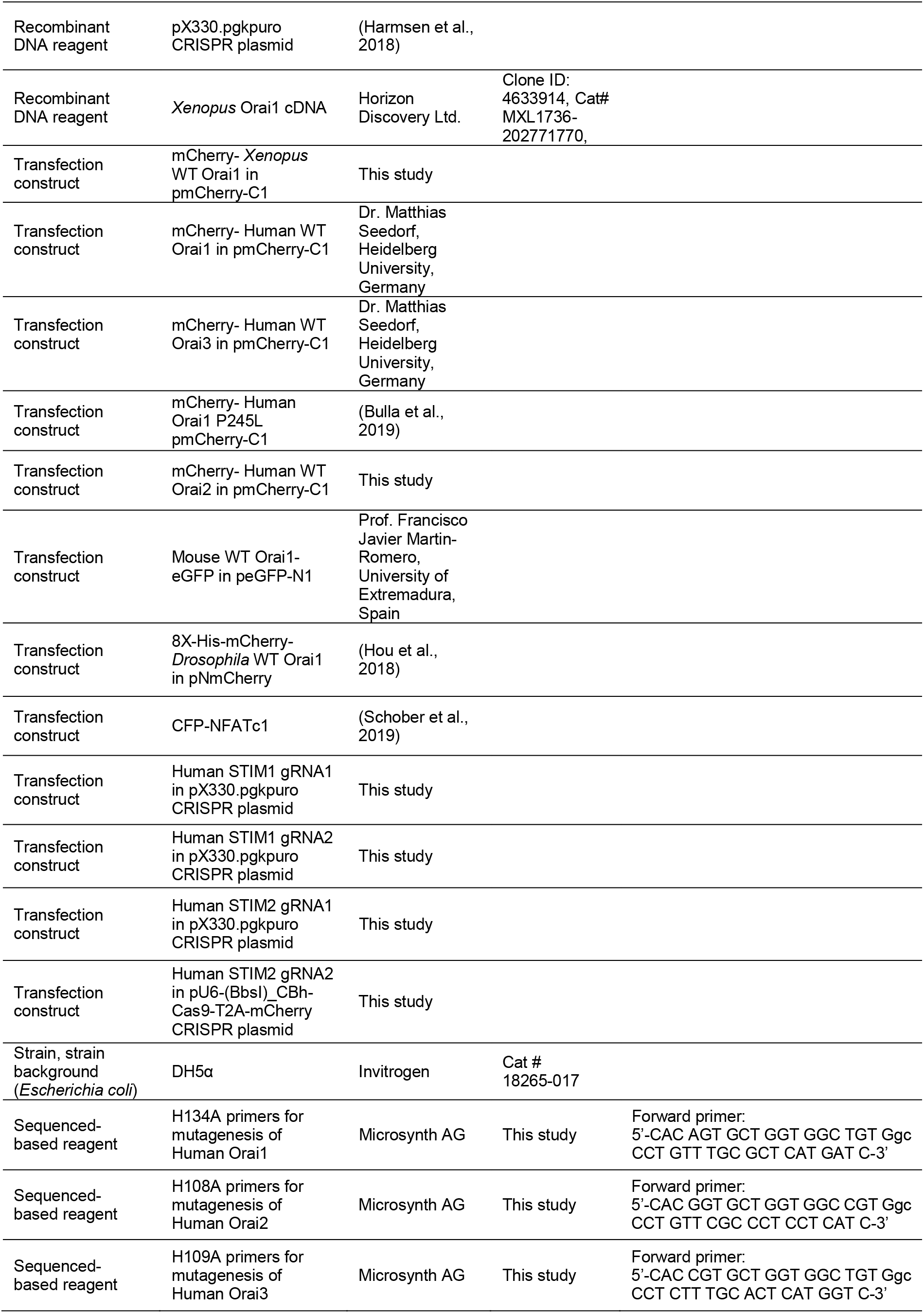
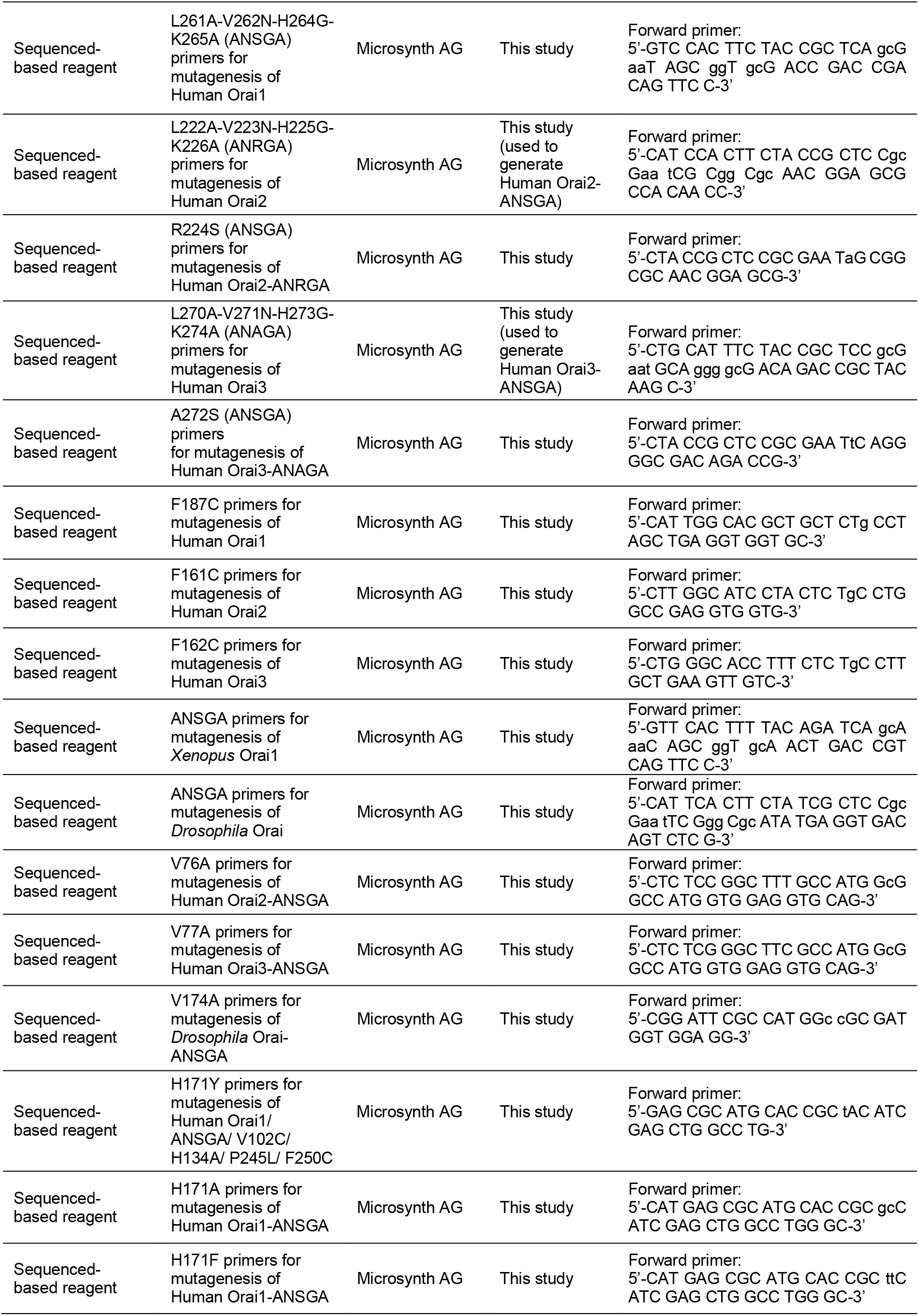
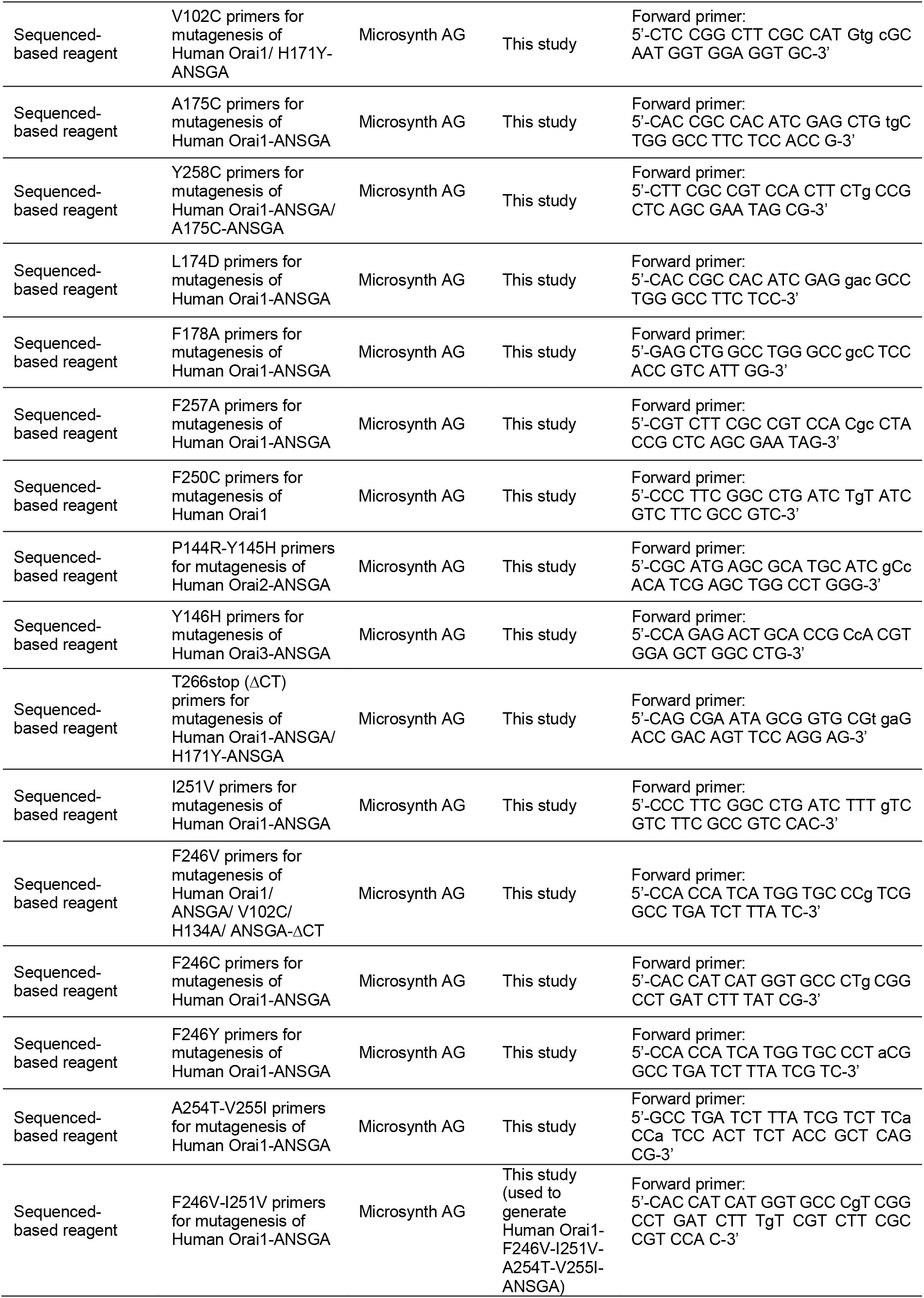
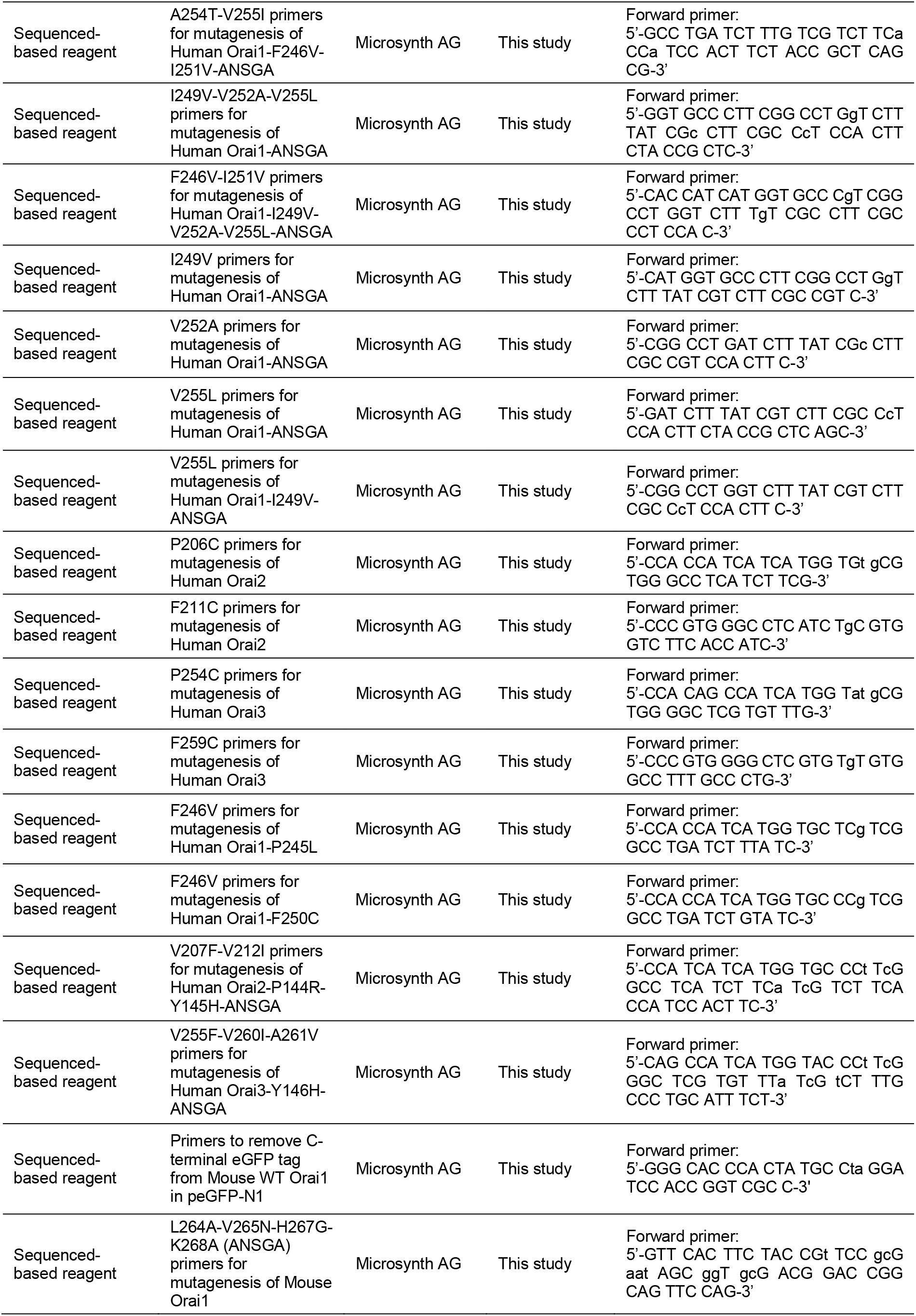
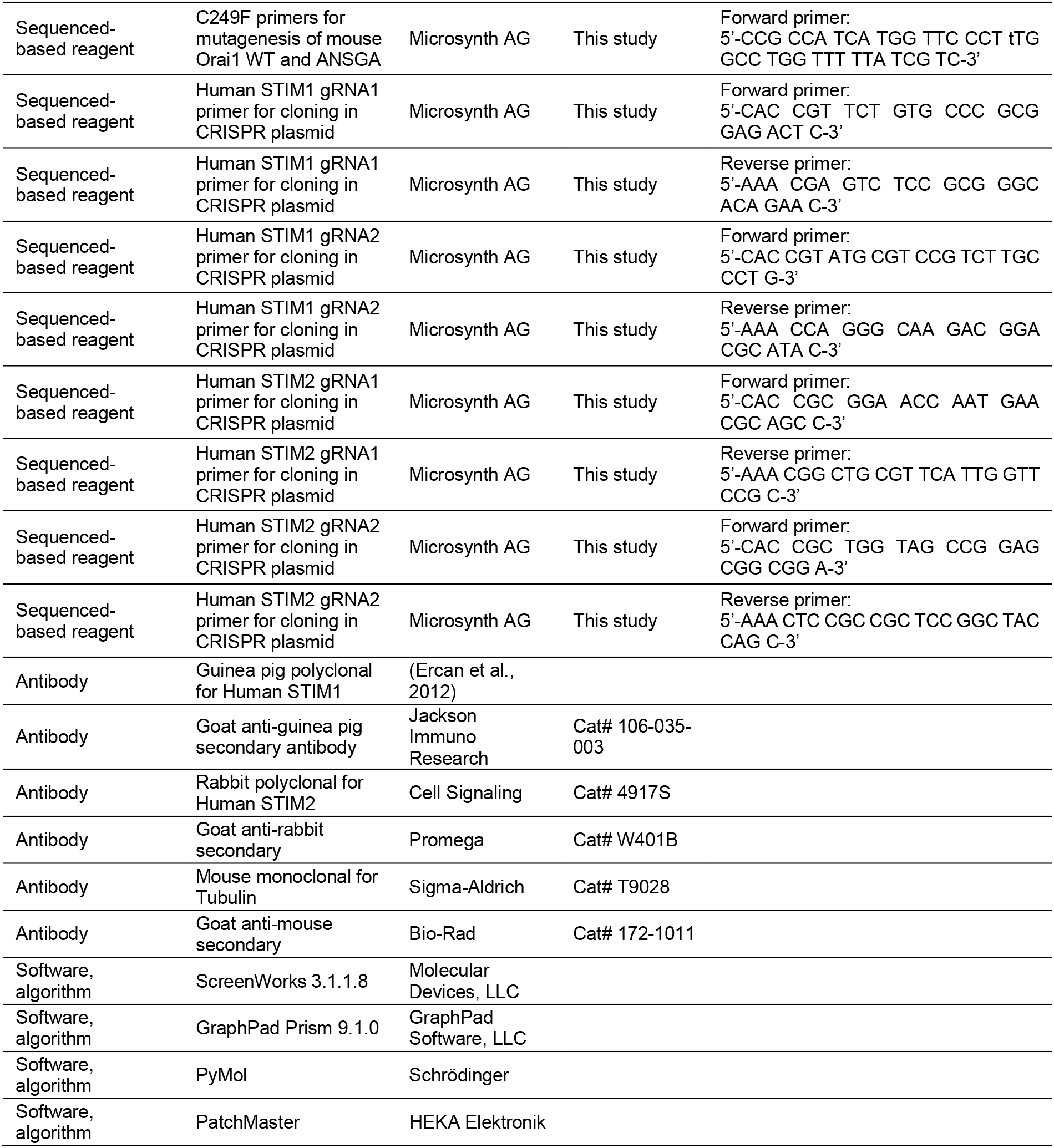

## Notes

### Competing Interest Statement

The authors have declared no competing interest.

